# Single Cell Transcriptomics of Fibrotic Lungs Unveils Aging-associated Alterations in Endothelial and Epithelial Cell Regeneration

**DOI:** 10.1101/2023.01.17.523179

**Authors:** Ahmed A. Raslan, Tho X. Pham, Jisu Lee, Jeongmin Hong, Jillian Schmottlach, Kristina Nicolas, Taha Dinc, Andreea M. Bujor, Nunzia Caporarello, Aude Thiriot, Ulrich H. von Andrian, Steven K. Huang, Roberto F. Nicosia, Maria Trojanowska, Xaralabos Varelas, Giovanni Ligresti

## Abstract

Lung regeneration deteriorates with aging leading to increased susceptibility to pathologic conditions, including fibrosis. Here, we investigated bleomycin-induced lung injury responses in young and aged mice at single-cell resolution to gain insights into the cellular and molecular contributions of aging to fibrosis. Analysis of 52,542 cells in young (8 weeks) and aged (72 weeks) mice identified 15 cellular clusters, many of which exhibited distinct injury responses that associated with age. We identified *Pdgfra^+^* alveolar fibroblasts as a major source of collagen expression following bleomycin challenge, with those from aged lungs exhibiting a more persistent activation compared to young ones. We also observed age-associated transcriptional abnormalities affecting lung progenitor cells, including ATII pneumocytes and general capillary (gCap) endothelial cells (ECs). Transcriptional analysis combined with lineage tracing identified a sub-population of gCap ECs marked by the expression of Tropomyosin Receptor Kinase B (TrkB) that appeared in bleomycin-injured lungs and accumulated with aging. This newly emerged TrkB^+^ EC population expressed common gCap EC markers but also exhibited a distinct gene expression signature associated with aberrant YAP/TAZ signaling, mitochondrial dysfunction, and hypoxia. Finally, we defined ACKR1^+^ venous ECs that exclusively emerged in injured lungs of aged animals and were closely associated with areas of collagen deposition and inflammation. Immunostaining and FACS analysis of human IPF lungs demonstrated that ACKR1^+^ venous ECs were dominant cells within the fibrotic regions and accumulated in areas of myofibroblast aggregation. Together, these data provide high-resolution insights into the impact of aging on lung cell adaptability to injury responses.

## Introduction

Lungs have a remarkable capacity to regenerate in response to injury, but this regenerative capacity deteriorates with aging resulting in disrepair and fibrosis (1–3). Pulmonary fibrosis, including idiopathic pulmonary fibrosis (IPF), is an aging-associated lung disorder characterized by alveolar damage and the accumulation of collagen-producing mesenchymal cells (myofibroblasts) into the lung interstitium resulting in excessive matrix accumulation (4, 5). IPF occurs largely in elderly adults, thus, age has emerged as a key risk factor (3, 6). Mechanisms linking aging to sustained fibroblast activation and the development of pulmonary fibrosis are not fully understood. Genomic instability, epigenetic alterations, metabolic dysfunction, cellular senescence, and stem cell exhaustion have been proposed as major contributors to the persistent fibrosis observed in aged individuals with IPF (1, 7–10).

By applying the bleomycin model of lung injury to aged mice, we and others have shown that these animals are more susceptible to develop sustained fibrogenesis after a single dose of bleomycin compared to young animals (9–13), suggesting that age-associated processes contribute to the persistent fibrosis that is observed in IPF. Impaired fibrosis resolution in the lungs of old mice was associated with elevated inflammation, senescence and enhanced collagen deposition relative to young mouse lungs (9, 11, 14, 15). Consistent with these observations, previous studies have shown that persistent fibrosis in the lungs of aged mice is characterized by the accumulation of senescent and apoptosis-resistant myofibroblasts, resulting in extended collagen secretion and impaired fibrosis resolution (15–17).

Fibrosis resolution is intrinsically and dynamically connected to regeneration (18, 19). Mechanisms driving lung regeneration following injury are not fully understood, and likely involve cross communications between different cell types in distinct lung anatomic locations. Loss of regenerative capacity in non-mesenchymal cells, such as epithelial and endothelial cells, has been proposed as an important contributory factor to the persistence of lung fibrosis in elderly patients with IPF (1, 11, 20). Lung progenitor stem cells, including epithelial and endothelial progenitors, have been recognized as important players in lung regeneration following injury (21–25), and several progenitors located in distinct locations throughout the lung have been described (21–24). For example, alveolar type-II epithelial (ATII) cells function as progenitor stem cells that give rise to alveolar type-I (ATI) epithelial cells following lung injury (22, 26). Recent advances in single-cell RNA sequencing (scRNA-seq) and cell lineage labeling have identified an alveolar differentiation intermediate population marked as *Krt8^+^* in mouse lungs post-bleomycin injury (27). This ATII-derived cell population was shown to transiently accumulate during alveolar regeneration, giving rise to ATI cells that replace the normal lung parenchyma (27). Notably, this intermediate population was found to be persistently present in pulmonary fibrosis patients (27), suggesting that aberrant epithelial cell differentiation may play an important role during disease progression.

Endothelial regeneration is also key to lung repair following injury, and this important function is orchestrated by lung endothelial progenitor cells known as general capillary (gCap) endothelial cells (ECs) (21, 23, 25). gCap ECs serve as specialized alveolar stem/progenitors that replace alveolar capillary ECs, including aerocytes (aCap ECs) which are often lost as a result of lung injury. We recently reported that the endothelial transcription factor ERG is a key orchestrator of gCap EC homeostasis and repair, and its dysfunctional signaling with aging impairs alveolar capillary regeneration, resulting in paracrine fibroblast activation and persistent fibrosis (9). The dynamics of epithelial and endothelial differentiation following lung injury are poorly understood, particularly with respect to how such cell populations are affected by aging. To gain insights into this unexplored feature of lung repair, we have carried out scRNA-seq on young and aged mouse lungs with and without injury induced by a single dose of bleomycin, focusing on cell population dynamics that associate with persistent lung fibrosis in aged animals. Our analyses revealed that impaired fibrosis resolution in aged mice following bleomycin challenge was accompanied by several transcriptional abnormalities largely affecting lung progenitor cells, including ATII and gCap ECs. We captured transcriptionally distinct ATII- and gCap EC-derived cell populations that exclusively appeared in bleomycin-treated lungs and likely represent activated intermediate states through which cells transit before their final commitment. We also discovered that the cellular composition and transcriptional signatures of these intermediate progenitors differed between young and aged lungs with those derived from aged lungs exhibiting several dysregulated signaling pathways, including those implicated in cellular metabolism, such as the YAP/TAZ, mTOR, and HIFα signaling pathways. These persistent aging-associated cellular states profoundly impacted lung homeostasis resulting in tissue hypoxia, disrepair, and sustained fibroblast activation. These findings suggest that metabolic maladaptation to injury with aging may interfere with the capacity of lung progenitors to produce functional and terminally differentiated cells following alveolar damage. Persistent accumulation of aberrant intermediate cells overtime may result in failure to regenerate lost lung parenchyma, continuous inflammation, and progressive fibrosis. Restoring regenerative programs in lung progenitor cells may facilitate their terminal differentiation and promote healthy healing of injured lung tissue.

Finally, we identified an expanding population of venous ECs, marked by the expression of *Ackr1*, in the lungs of fibrotic aged mice which was topologically scar-restricted and exhibited angiogenic and inflammatory features. Thus, aging may also alter the endothelial composition of the lung microvasculature promoting the accumulation of a venous-type pro-inflammatory phenotype. Our observations offer a picture of the cellular dynamics associated with sustained fibrosis in aged lungs and suggest that maladaptation to injury with aging may interfere with the capacity of lung progenitors to produce functional and terminally differentiated cells thereby affecting fibrosis resolution and restoration of normal lung functions.

## Results

### Single-cell atlas of young and aged mouse lungs during fibrosis resolution

To define age-associated cellular and transcriptional responses to lung injury-induced fibrosis, we employed scRNA-seq using the 10X Genomic platform to profile the whole lung digests isolated from young mice (2 months) and aged mice (18 months) following intratracheal bleomycin instillation. Young and aged lungs were isolated at the early resolution phase of fibrosis (30 days post-bleomycin) (Fig. 1A) and were compared to saline-treated young and aged lungs. After quality filtering, we obtained approximately 52,542 cell profiles, from all the samples. We then performed dimensionality reduction with canonical correlation analysis (CCA) subspace alignment followed by unsupervised clustering (Fig.1B). Using canonical lineage-defining markers to annotate clusters, we identified 15 distinct lung cell types (Fig.1B and Supplement Fig.1). Whole cluster analysis demonstrated that pulmonary endothelial cells were the most abundant cell type that was captured and revealed two distinct endothelial cell clusters: Endothelial I (42.82%) which express gCap EC markers (*Gpihpb1*, *Plvap*, *Tek*), and Endothelial II (4.3%) which is enriched in aCap markers (*Car4*, *Ednrb*, *Igfbp7*) (Fig.1B and Supplement Fig.1). Additional analysis using representative marker genes identified clusters of endothelial, epithelial, and mesenchymal cells, as well as populations of immune cells, including macrophages, monocytes, neutrophils, and natural killer cells (Fig.1C). Bleomycin-induced lung injury caused drastic alterations mainly in endothelial, epithelial, and mesenchymal cells, as evidenced by the appearance of additional distinct cell clusters derived from these lineages in the lungs of young and aged mice following bleomycin injury (Fig.1D).

**Figure 1.**
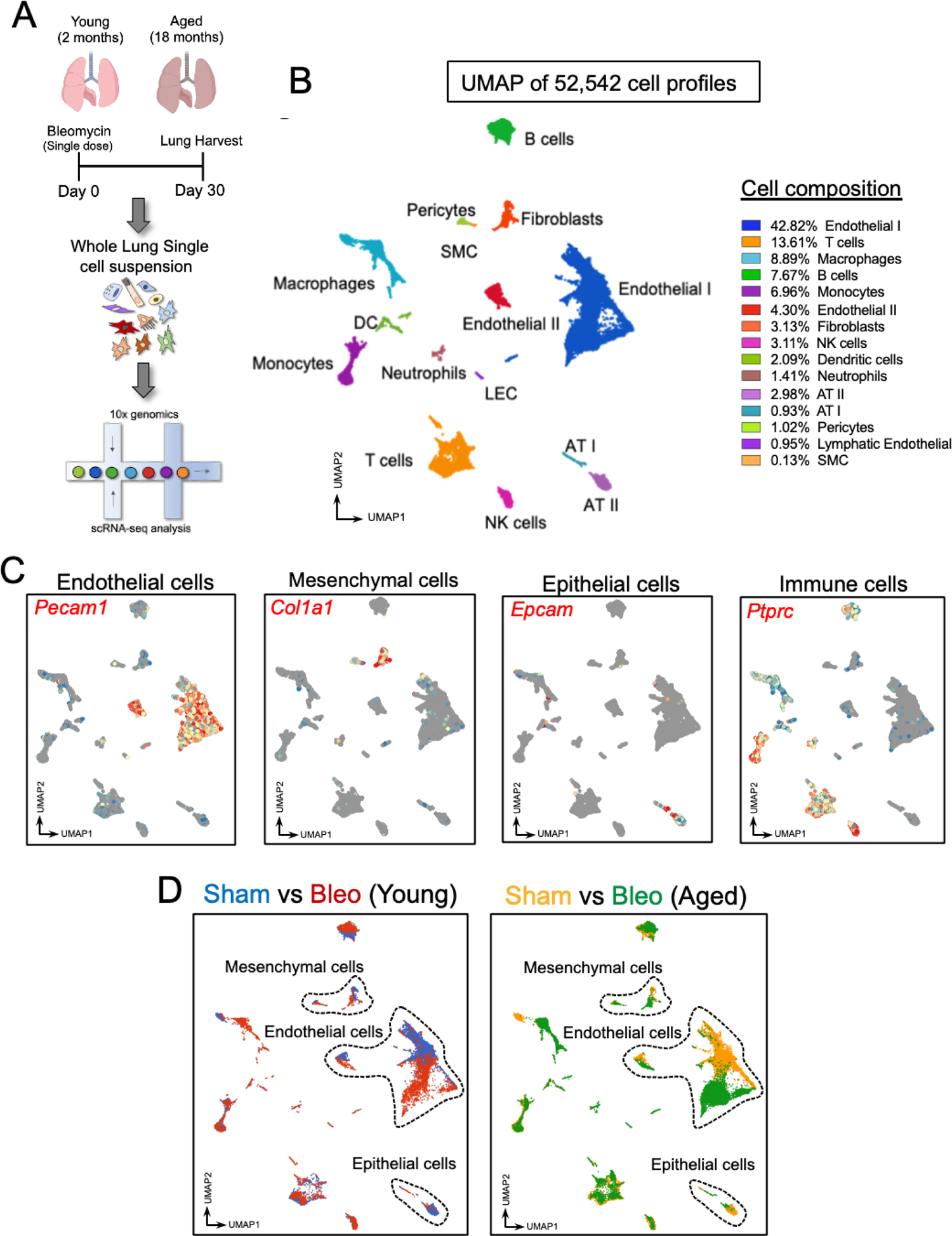
scRNA-seq analysis of young and aged mouse lungs following bleomycin challenge. (**A**) Young sham (n=1), aged sham (n=2), bleomycin-treated young (n=1), and bleomycin-treated aged (n=2) lungs were harvested at day 30 post bleomycin deliver and prepared for scRNA-Seq analysis. **(B)** UMAP embedded visualization of the cells from all lung samples shows different cell populations and their composition. **(C)** UMAP plots of single cells showing gene expression for indicated marker genes. **(D)** UMAP shows different cell clusters from untreated and bleomycin-treated young and aged lungs. Colored dots highlight endothelial, epithelial, and mesenchymal cluster shifts in bleomycin-treated young and aged lungs relative to untreated ones.

### Alveolar fibroblast activation following injury is enhanced in aged lungs

To characterize the cellular and transcriptional responses of lung mesenchymal cells, which are the major contributors to interstitial collagen production upon bleomycin injury (4, 5), we re-clustered mesenchymal cell lineages and identified 6 distinct cell populations (Fig.2A,B). One of these mesenchymal cell clusters emerged exclusively in bleomycin-treated lungs co-expressing markers of adventitial and alveolar fibroblasts, including *Pdgfra*, *Cdh11*, and *Mfap4* and markers of activated fibroblasts (myofibroblasts), such as *Tnc, Fn1, Timp1, and Ctsk* (Fig.2C). Of note, the appearance of a population of activated fibroblasts in bleomycin-treated lungs was accompanied by the reduction of quiescent alveolar fibroblasts, suggesting that alveolar fibroblasts are the major source of myofibroblasts upon bleomycin injury (Fig.2D,E). Volcano plots shown in Fig.2F identified upregulated and downregulated genes in injured young and aged lung fibroblasts. Among the upregulated genes were numerous pro-fibrotic genes implicated in ECM remodeling, including *Col3a1*, *Timp1*, *Igfbp7*, and *Fn1*. In depth gene expression analysis revealed that numerous fibrotic genes, including *Spp1*, *Acta2*, *Ldha*, and *Timp1*, exhibited increased expression in activated fibroblasts from aged lungs relative to those from young ones (Fig.2G), consistent with our prior bulk RNA-seq studies that demonstrated that lung fibroblasts in aged mice fail to undergo quiescence and sustain ECM protein deposition following bleomycin-injury (10).

**Figure 2.**
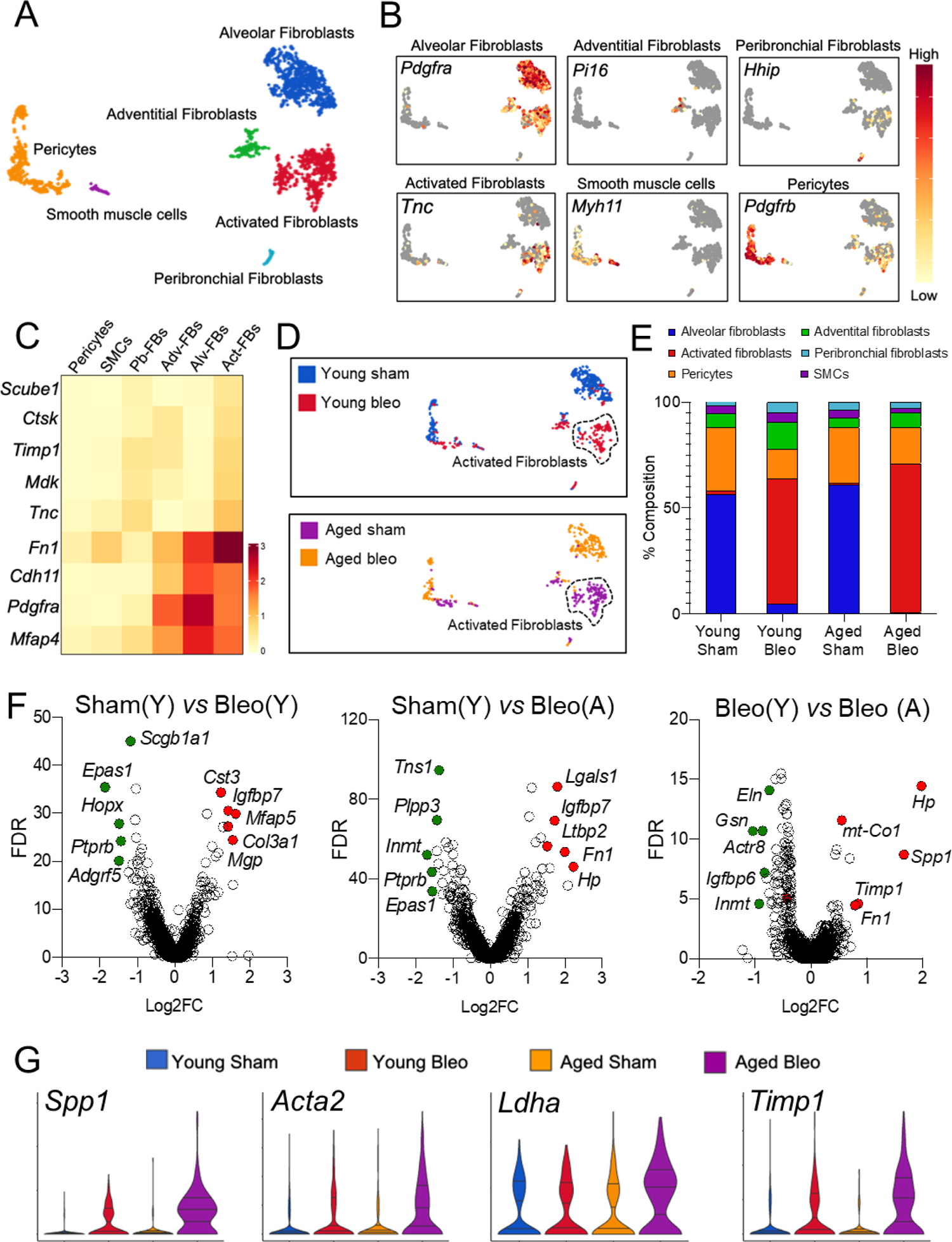
Mesenchymal cell responses in fibrotic young and aged mouse lungs. **(A)** UMAP embedded visualizations of six different mesenchymal cell clusters. **(B)** UMAP plots exhibiting distinctive marker genes defining different lung mesenchymal lineages. **(C)** Heatmaps showing differentially expressed marker genes in each mesenchymal cell clusters**. (D)** UMAP plots displaying mesenchymal cell clusters in young (Blue) and aged (Orange) uninjured lungs and young (Red) and aged (Purple) injured lungs. Circled are activated fibroblasts from bleomycin-treated young and aged lungs. **(E)** Percentage of mesenchymal cell subsets grouped by age and treatment**. (F)** Volcano plots showing the distribution of differentially expressed genes in different comparisons based on fold changes and significance. **(G)** Violin plots showing increased expression of pro-fibrotic genes in injured aged lungs relative to other conditions.

### Alveolar epithelial intermediate state composition is altered in fibrotic aged lungs and associated with impaired YAP/TAZ signaling

Alveolar epithelium plays an important role during lung homeostasis and repair following injury (22, 28, 29). ATII cells exhibit critical secretory functions and regenerative roles in the alveolus; however, their aberrant turnover has been observed in fibrotic lungs and contributes to the development of IPF (30–32). To characterize the heterogeneity and injury responses of alveolar epithelial cells in aging, we focused on Epcam^+^ epithelial cells that expressed high levels of ATII and ATI markers. Re-clustering of Epcam^+^ alveolar epithelial cells identified three distinct subpopulations: ATII, which was the most abundant alveolar epithelial cell cluster, ATI, and an intermediated cluster of alveolar epithelial cells (here defined as intAE) that co-expressed both ATII and ATI markers (Fig.3A). Notably, this intermediate cell population appeared almost exclusively in bleomycin treated lungs (young and aged) and was accompanied by the reduction of ATII progenitor cells, suggesting that this intermediate cell population emerged from ATII cells as they differentiate into ATI cells (Fig.3B,C). While this intermediate cell cluster shared numerous markers with ATI cells, such as *Hopx*, *Vegfa*, and *Rtkn2*, it also expressed a distinct set of genes, including *Clu*, *Krt8*, and *Cdkn1a* that were weakly or not expressed by differentiated ATI or ATII cells (Fig.3D). These findings are consistent with a previous study demonstrating the appearance of intermediate Krt8^+^ ATII-derived cells in the lungs of mice treated with bleomycin (27).

**Figure 3.**
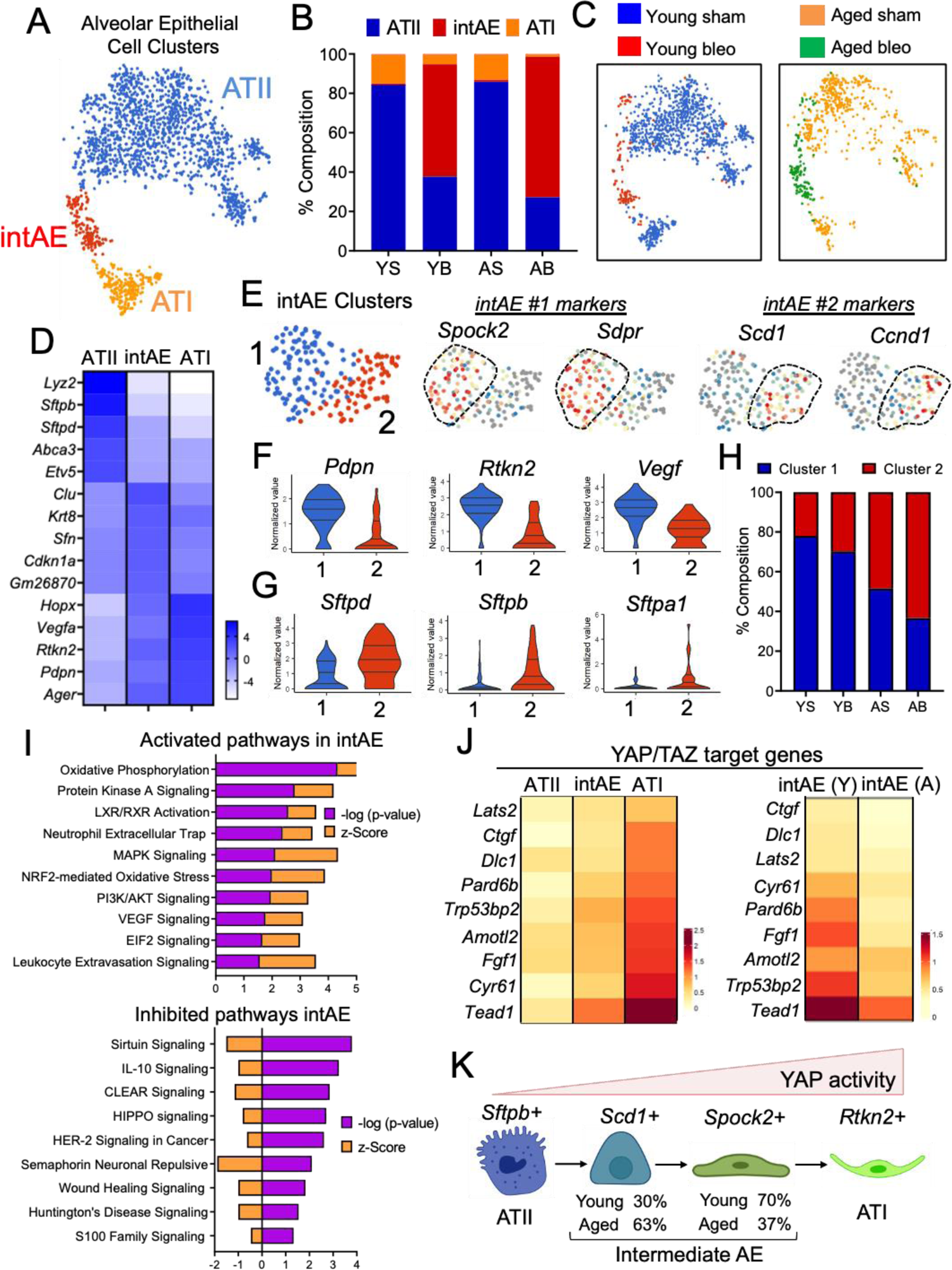
Aberrant alveolar epithelial cell differentiation in fibrotic mouse lungs with aging. **(A)** UMAP plot showing different alveolar epithelial cell clusters. ATII (Blue), intermediate AE (intAE, Red), and ATI (Orange). **(B)** Percentage of alveolar epithelial subsets grouped by age and injury. **(C)** UMAP plots displaying alveolar epithelial cell clusters in young (Blue) and aged (Orange) uninjured lungs, and young (Red) and aged (Green) injured lungs. **(D)** Heatmap showing the average expression level of marker genes across different alveolar epithelial cell clusters. **(E)** UMAP visualization of intermediated alveolar epithelial cell clusters and marker gene signatures. **(F)** Violin plots showing the expression of ATI cell marker genes in intAE clusters. **(G)** Violin plots showing the expression of ATII cell marker genes in intAE clusters. **(H)** Cell composition in each intAE cluster. Most intermediate alveolar epithelial cells exhibiting ATII marker genes (less differentiated) were from bleomycin-treaded aged lungs. **(I)** Ingenuity pathway analysis shows canonical pathways enriched in aged lung intAE cells relative to young ones. *P* values were generated in IPA using Fisher’s test (log2 FC ≤-0.1 or ≥0.1, *p* value ≤0.05). *P* value and activation z-score were used for plotting canonical pathways and activated upstream regulators respectively. **(J)** Heatmaps showing the average expression levels of YAP/TAZ target genes in different alveolar epithelial cell clusters and in aged intAE versus young intAE**. (K)** Schematic showing the YAP-dependent transition of ATII into ATI cells and the accumulation of undifferentiated states with aging.

To gain further insights into the impact of aging on the intermediate cell population we re-clustered *Krt8*^+^ *Cdkn1a*^+^ epithelial cells and obtained two distinct sub-clusters (Intermediate Cluster 1 and Intermediate Cluster 2) (Fig.3E), with each sub-cluster expressing a unique set of genes, including *Spock2* and *Sdpr* in cluster 1 and *Scd1* and *Ccnd1* in cluster 2 (Fig.3E). Differential gene expression analysis revealed that intermediate cluster 1 was enriched in cells expressing ATI markers, such as *Pdpn*, *Rtkn2*, and *Vegf*, (Fig.3F), whereas cells in the intermediate cluster 2 expressed high levels of ATII markers, including *Sftpd*, *Sftpb*, and *S*ftpa1 (Fig.3G). Intriguingly, we found that cluster 1 (high ATI markers) was largely occupied by cells derived from uninjured and injured young lungs, whereas cluster 2 (high ATII markers) was occupied primarily by cells derived from uninjured and injured aged lungs (Fig.3H). These findings suggested that following lung injury ATII cells pass through multiple intermediate transcriptional states before transitioning into fully functional ATI cells and suggested that these differentiation steps are impaired with aging resulting in aberrant ATI cell differentiation. Because intermediate ATII cells were reported to accumulate in the lungs of elderly patients with IPF (27), our findings strongly suggest that ATII to ATI differentiation is impaired with aging, resulting in the accumulation of epithelial intermediates that poorly expressed ATI markers and may potentially have deleterious effects on lung fibrosis resolution.

To investigate signaling pathways whose dysregulation may result in aberrant ATII differentiation with aging, we compared the transcriptional signatures of intermediate alveolar epithelial cells between young and aged lungs, followed by gene ontology (GO) analysis. As shown in Fig.3I, we identified numerous activated pathways in aged intermediate alveolar epithelial (intAE) cells relative to young ones, and among them were those implicated in oxidative phosphorylation, oxidative stress, cell proliferation, as well as cell survival, such as MAPK and PI3K/AKT signaling pathways. Among the inhibited signaling pathways in aged intAE cells were those previously implicated in cellular rejuvenation, wound healing, and metabolism, including, Sirtuin, CLEAR, and Hippo signaling pathways. Intriguingly, the Hippo pathway and its downstream effectors YAP and TAZ have been recently shown to play a key role during the differentiation of ATII cells into ATI (33–35), and dysfunctional activation of YAP/TAZ signaling pathway has been associated with advanced aging (36). These studies prompted us to compare the expression of YAP/TAZ target genes in alveolar epithelial cells during their transition to mature ATI cells, which revealed that several YAP/TAZ target genes showed increased expression during the transition of ATII into ATI cells, with ATI cells exhibiting the strongest expression of YAP/TAZ target genes (Fig.3J). However, compared to young lungs, aged intAE cells exhibited reduced expression of YAP/TAZ target genes, suggesting that impaired YAP/TAZ activity may contribute to defective alveolar epithelial cell differentiation, leading to the accumulation of intermediate states resulting in disrupted lung repair (Fig.3K).

### Lung injury induces capillary endothelial cell transition to undifferentiated states that persist in aged and IPF lungs

We recently demonstrated that animal aging impairs pulmonary vasculature functions during lung repair and fibrosis resolution contributing to capillary loss and sustained fibrosis (9, 11). To investigate endothelial cell heterogeneity and transcriptional responses to injury with aging, we re-clustered lung endothelial cells (ECs) (*Cldn5^+^* and *Pecam-1^+^)* using unsupervised clustering analysis. To annotate endothelial subpopulations, we used lineage-defining markers based on prior studies and identified 9 endothelial cell subclusters, including those from veins, arteries, lymphatics, gCap, aCap, proliferating ECs, and endothelial tip cells (*Hspg2*^+^) (Fig.4A and Supplement Fig.2). Other endothelial cell subclusters also included, activated gCap ECs and activated venous ECs, which were only apparent in bleomycin-injured lungs (young and aged) (Fig.4A and Supplement Fig.2). Although these newly emerged activated EC subpopulations exhibited distinct gene expression signatures defining their different endothelial origin, they also shared common markers of cell activation, including *Ankrd37*, *Ntrk2*, *Cxcl12,* and *Tmem252* (Supplement Fig. 2), which can distinguish them from their quiescent counterparts. These findings suggest that, similarly to airway and alveolar epithelial cells, gCap and venous ECs proceed through common intermediate states in response to injury before their final commitment, and their transient appearance may be relevant to the vascular repair process. Given their plasticity and implications in lung injury and fibrosis, we chose to focus follow-up studies on endothelial cells derived from gCap and venous ECs.

**Figure 4.**
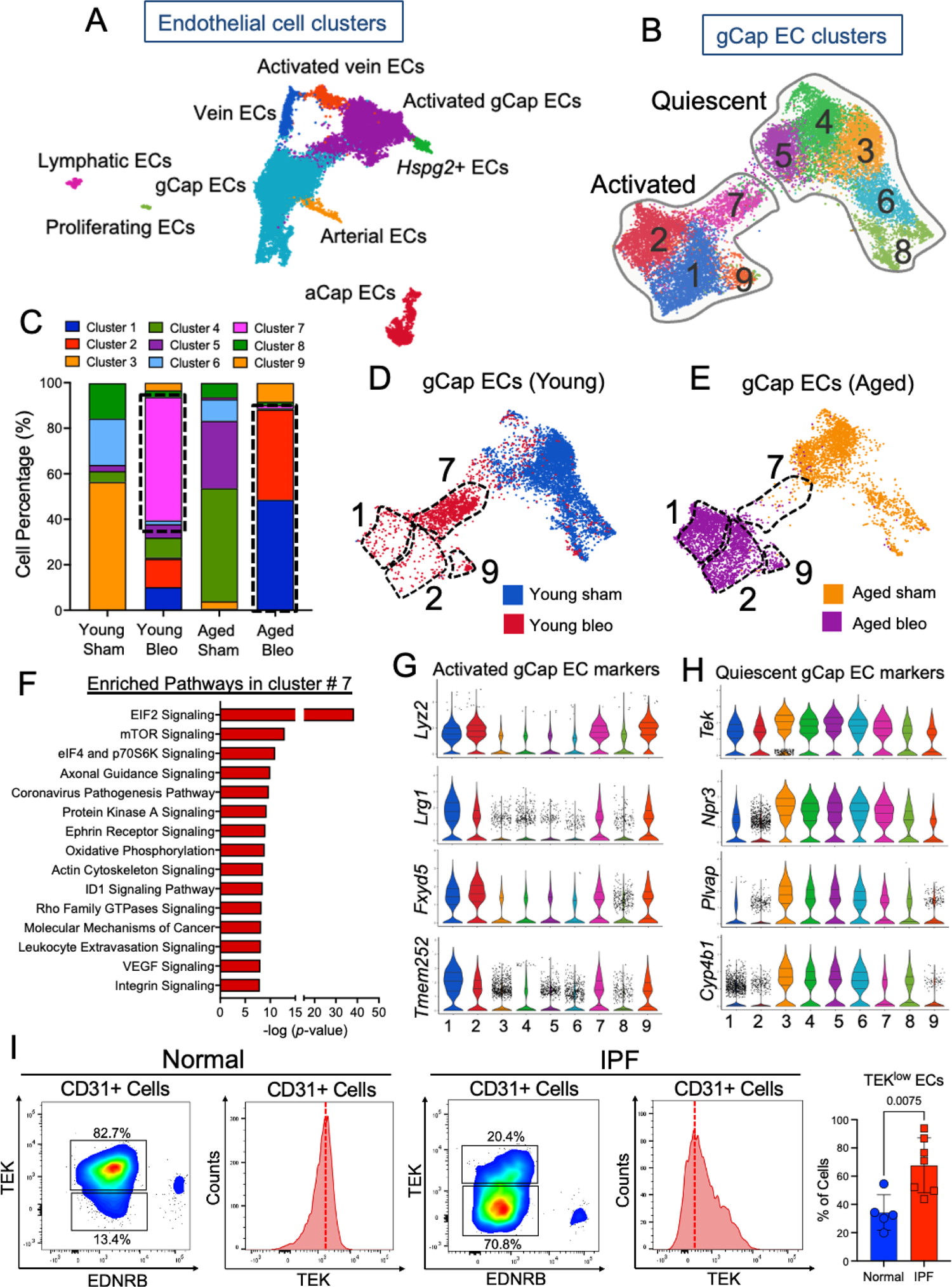
Lung injury triggers gCap EC transition to undifferentiated states that persisted in aged and IPF lungs. **(A)** UMAP plot of all endothelial cells (ECs). Each color indicates a distinct EC state at baseline and in response to injury. **(B)** UMAP plot of gCap EC subtype. Cells in clusters 1,2, 7, and 9 are activated gCap ECs that emerged exclusively in bleomycin-treated lungs. **(C)** Composition of gCap EC clusters across different groups. **(D, E)** UMAP plots displaying gCap EC clusters in young (Blue) and aged (Orange) uninjured lungs, and young (Red) and aged (Purple) injured lungs. Clusters 1, 2 and 9 were largely occupied by injured aged gCap ECs whereas cluster 7 by injured young gCap ECs. **(F)** Ingenuity pathway analysis shows canonical pathways enriched in gCap EC cluster #7 relative to other gCap EC clusters. *P*-values were generated in IPA using Fisher’s test (log2 FC ≤-0.1 or ≥0.1, *p* value ≤0.05). *P* value and activation z-score were used for plotting canonical pathways and activated upstream regulators respectively. **(G)** Violin plots showing the *de novo* expression of activated EC marker genes across different gCap EC clusters. **(H)** Violin plots showing the expression of quiescent gCAP EC marker genes across different clusters. Activated gCap EC (1, 2, 9) exhibit the strongest reduction of canonical gCap EC marker genes **(I)** FACS analysis of normal and IPF lungs. Antibodies against TEK and EDNRB were used to discriminate gCap EC from aCap EC. Out of all CD31 positive cells gCap ECs were defined as (TEK^High^ EDNRB^Low^) whereas aCap ECs strongly expressed EDNRB and were largely negative for TEK. IPF lungs exhibited an increased percentage of ECs expressing low levels of the gCap marker TEK compared to healthy lungs. Values are summarized as mean and SD and analyzed using a two-tailed Student’s *t*-test. (Normal n = 5; IPF n = 7).

As lung gCap ECs exhibit stemness properties (21, 23), and give rise to new capillary ECs, including aCap, following injury (21, 23), we first investigated whether gCap EC turnover and transcriptional responses to injury were compromised with aging. Cluster analysis using previously discovered gCap EC markers revealed 9 distinct clusters of capillary ECs across all groups (Fig.4B). These clusters can be further separated into quiescent gCap ECs (3, 4, 5, 6, 8), which are predominant in the sham groups (young and aged), and activated gCap ECs (1, 2, 7, 9), which emerged in the bleomycin-treated groups only (young and aged). Among the quiescent capillary ECs, those in clusters 3 and 6 were abundant in the young sham group whereas those in clusters 4 and 5 largely appeared in the sham aged group, suggesting baseline differences in the transcriptional signatures of young and aged gCap ECs. Consistent with these findings, we previously demonstrated that in absence of injury, lung ECs from aged mice were epigenetically distinct from young ones (9), emphasizing their baseline transcriptional differences.

Capillary ECs in clusters 1, 2, 9, and 7 (activated gCap ECs) exclusively surfaced in bleomycin-treated groups (Fig.4C). Intriguingly, while these activated EC emerged in both young and aged lungs post injury, their cellular composition and transcriptional signatures varied considerably with aging. For example, cluster 7 appeared exclusively in the injured young gCap EC group and was the most abundant cluster relative to other capillary clusters in the young group (Fig.4D,E). IPA pathway analysis of highly expressed genes in cluster 7 identified numerous enriched pathways previously implicated in vascular remodeling and regeneration, including mTOR, Ephrin Receptor, Axonal Guidance, VEGF, and Integrin pathways (Fig.4F). While ECs in cluster 7 share some gene markers with those in clusters 1, 2, and 9, including the activated cell markers *Lyz2*, *Lrg1*, *Fxyd5*, and *Tmem252* (Fig.4G), they were overall transcriptionally more similar to quiescent gCap ECs, as evidenced by the elevated expression of canonical gCap EC markers, such as *Tek*, *Npr3*, and *Plvap*, which was strongly attenuated in activated gCap ECs in clusters 1, 2 and 9 (Fig.4H). These findings demonstrated that cluster 7 arises in the lung of young mice during lung fibrosis resolution and is largely absent in fibrotic aged lungs, suggesting that this newly formed endothelial state may play a role in restoring a functional capillary bed, and this regenerative function is lost with aging.

Clusters 1 and 2 largely emerged in the injured aged group and were markedly characterized by the increased expression of endothelial cell activation markers and by the reduced expression of gCap EC differentiation markers (Fig.4G,H), suggesting that injury in aged lungs resulted in the accumulation of undifferentiated, possibly dysfunctional, capillary ECs that may perpetuate injury responses and fibrosis. Intriguingly, gene expression analysis of clusters 1, 2 and 7 showed that cells in these clusters are transcriptionally coupled, and they represent three distinct but connected intermediate states (Fig. Supplement 3). Given that cells in cluster 7 appear almost exclusively in young lungs during fibrosis resolution (Fig.4C-E), these findings suggest that injured aged gCap ECs in cluster 1 and 2 are unable to terminally differentiate into mature gCap or aCap ECs resulting in their accumulation.

To extend these findings to human disease and investigate the capillary EC turnover in the context of progressive lung fibrogenesis, we carried out a fluorescence activated cell sorting (FACS) analysis on diseased lungs from patients with idiopathic pulmonary fibrosis (IPF). We developed a FACS strategy using antibodies against markers that were enriched in capillary ECs relative to ECs from other vascular beds, including TEK (gCap ECs), EDNRB (aCap ECs), and CD31 (pan-ECs marker). The overall number of CD31-expressing cells was lower in IPF lungs compared to healthy ones (data not shown). Furthermore, we discovered that within each CD31^+^ EC population, the number of undifferentiated capillary ECs (CD31^+^ TEK^Low^) was significantly elevated in IPF lungs compared to healthy ones (Fig.4I), suggesting that, similarly to fibrotic aged mouse lungs, human fibrotic lungs exhibited increased number of intermediate capillary ECs relative to healthy lungs resulting in aberrant capillary EC turnover.

### Increased YAP activation, glycolysis and hypoxia are associated with impaired gCap EC differentiation in fibrotic aged lungs

Our clustering analysis of EC transcriptomes showed that during the early phase of lung fibrosis resolution the composition of gCap ECs varies substantially between young and aged mice, with aged mice exhibiting an increased number of activated/undifferentiated gCap ECs relative to young mice, suggesting impaired gCap EC maturation following injury in aged mice. To shed some light on these dysfunctional capillary dynamics we carried out a gene enrichment analysis to identify aging-associated gene signatures in these activated EC populations that can serve as indicators or their altered cellular state. Since clusters 1 and 2 were the most abundant clusters of undifferentiated gCap ECs (Fig.4C), we focused our analysis on these two endothelial subtypes. Volcano plot in Fig.5A shows upregulated and downregulated genes in clusters 1 and 2 relative to all other capillary EC clusters. Violin plots displayed in Fig.5B highlights enriched genes in capillary ECs from clusters 1 and 2, which include canonical HIF1α- and hypoxia-regulated genes, such as *Ankrd37*, *Bnip3,* and *Bhlhe40*. Of note *Bnip3* encodes for a BH3 protein which plays a key role in HIF1α-regulated mitochondrial bioenergetics and cell death in response to injury or during hypoxic conditions (37–40). Dysfunctional BNIP3 signaling in aging resulted in reduced mitophagy (the removal of damaged mitochondria) (41–43), impaired cellular metabolism (44), and elevated inflammation (43), suggesting that aberrant lung capillary EC turnover in aged mice may be the direct result of an abnormally prolonged activation of HIF1α signaling resulting in metabolic maladaptation to injury. HIF1α signaling was previously shown to regulate numerous genes encoding for glycolytic mediators (45, 46), and HIF1α-stimulated glycolysis is key to satisfy energetic requirements during cell reprogramming following injury, hypoxic stress, or when mitochondrial respiration is compromised (45). Consistent with this adaptive mechanism, we found that gCap ECs in clusters 1 and 2 expressed high levels of glycolytic genes such as *Eno1*, *Pfkl*, *Aldoa* and *Ldha* (Fig.5C). Furthermore, IPA and upstream regulators analysis identified the enrichment of numerous signaling regulators linked to metabolic control, including mTOR, MYC, YAP1, and HIF1α, that were associated with the appearance of cluster 1 and 2 (Fig.5D,E), suggesting that dysregulated metabolic dynamics in aged gCap ECs underlie their abnormal turnover in response to injury. Among the top upstream regulators that emerged in clusters 1 and 2 (Fig.5E), was the Hippo pathway effector YAP. Aside for its known function as mechanoregulator, YAP and its paralog TAZ have been implicated in metabolic reprogramming during cell growth and differentiation (47–49). YAP/TAZ-mediated transcription has been shown to regulate varying metabolic pathways, including glycolysis, mitochondrial respiration, and autophagy (50–53), which are required for cellular adaption to adverse environmental conditions, such as injury or oxygen deprivation.

**Figure 5.**
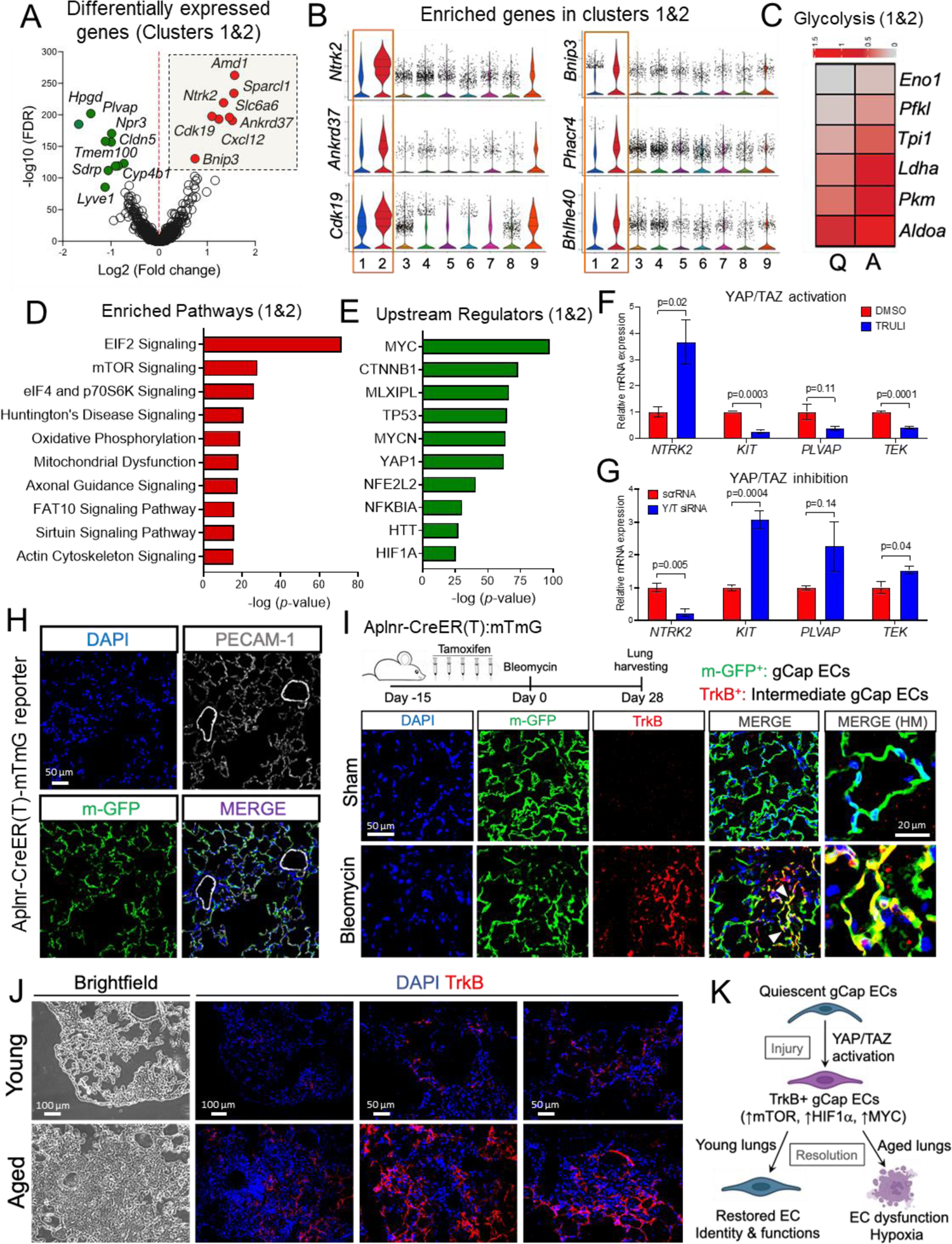
Augmented metabolic pathways and hypoxia are associated with aberrant gCap EC differentiation in fibrotic aged lungs. **(A)** Volcano plots showing the distribution of differentially expressed genes in gCap EC clusters 1 and 2 compared to other gCap EC clusters. **(B)** Violin plots showing the expression genes enriched in cluster 1 and 2. **(C)** Heatmap showing average expression of glycolysis genes in quiescent (Q) versus activated (A) gCap EC clusters. **(D, E)** Ingenuity pathway analysis shows canonical pathways and upstream regulators enriched in clusters 1 and 2 relative to other gCap EC clusters. *P* values were generated in IPA using Fisher’s test (log2 FC ≤-0.1 or ≥0.1, *p* value ≤0.05). *P* value and activation z-score were used for plotting canonical pathways and activated upstream regulators respectively. **(F, G)** qPCR analyses of human lung microvascular ECs (HLMECs) treated with the LATS1/2 inhibitor TRULI and the siRNAs targeting YAP and TAZ for 48 hours. YAP activation in these cells partially recapitulates the gene expression signature observed in activated gCap ECs. Values are summarized as mean and SD and analyzed using a two-tailed Student’s *t*-test. (N = 3). **(H)** Immunofluorescence images showing genetically labeled gCap ECs (membrane-GFP) in uninjured lungs of Aplnr-CreER(T)-mTmG reporter mouse. Lungs from these mice were harvested ten days following the last dose of tamoxifen (five total doses). An antibody against the pan-endothelial cell marker PECAM-1 was used to visualize all lung ECs. PECAM-1 positive ECs from large vessels showed no GFP expression. **(I)** Schematic representation of the approach used to detect TrkB-expressing gCap ECs following bleomycin injury. gCap ECs (mGFP) were lineage labeled in Aplnr-CreER(T)-mTmG mice 15 days prior to bleomycin administration (Day 0). Sham and bleomycin-injured lungs were harvested 28 days post bleomycin delivery and subjected to immunofluorescence analysis. An antibody against TrkB was used to detect injured gCap ECs (Red). gCap ECs co-expressing GFP and TrkB (yellow) only emerged in injured lungs (n=2). **(J).** Immunofluorescence images showing TrkB-expressing cells in the lungs of young and aged mice 37 days post bleomycin challenge (n=2). **(K)** Schematic showing putative mechanisms implicated in of gCap EC remodeling in response to lung injury and during lung fibrosis resolution.

To study the contribution of YAP/TAZ activation or inhibition to capillary EC behavior, we treated human lung microvascular ECs (HLMVECs) with TRULI, a recently discovered small-molecule inhibitor of LATS kinases that promotes YAP/TAZ activation (54). HLMVECs treated with this inhibitor exhibited increased expression of *NTRK2,* a marker of gCap EC activation, and reduced expression of the canonical capillary markers *TEK*, *KIT*, and *PLVAP* (Fig.5F). Inversely, YAP and TAZ knockdown in these cells potently inhibited *NTRK2* expression and upregulated *TEK*, *KIT* and *PLVAP* gene expression (Fig.5G). These findings together with our pathway analysis support a role for YAP/TAZ in capillary EC transition to an activated state following lung injury before returning to quiescence or giving rise to mature aCap ECs. As intermediate gCap ECs largely accumulated in the lungs of aged mice with delayed fibrosis resolution, these findings suggest that persistent YAP/TAZ signaling in aged gCap ECs contributes to their aberrant differentiation, resulting in disrupted lung homeostasis and progressing scarring following injury. Given that YAP/TAZ signaling pathway cooperates with several metabolic regulators including HIFα and MYC (50, 52, 53, 55), to regulate injury responses, we reasoned that a dysregulated crosstalk between these metabolic regulators in aged gCap ECs may underly their aberrant behavior.

To shed further insights into gCap EC activation following lung injury and substantiate our transcriptional analysis, we carried out a lineage tracing experiment using a conditional Aplnr-Cre-ER(T): *Rosa26-mdtTomato/mGFP* reporter mouse (Aplnr-Cre-ER(T)-mTmG), in which gCap ECs are permanently labeled with GFP upon tamoxifen administration. Lineage labeled gCap ECs (mGFP) were exclusively located in the lung alveoli and expressed the pan-endothelial cell marker PECAM-1 (Fig.5H). Notably, PECAM-1 positive ECs from large vessels, such as veins and arteries, were not genetically labeled (Fig.5H). Bleomycin was intratracheally delivered to the lungs of these mice and the expression of the intermediate gCap marker and YAP/TAZ-regulated target tropomyosin receptor kinase B (TrkB, encoded by *Ntrk2*) was assessed at 28 days post bleomycin administration (Fig.5I). Immunofluorescence analysis showed that TrkB expression was acquired exclusively in gCap ECs of injured lungs and was undetectable in those from uninjured lungs (Fig.5I).

To assess the impact of aging on the appearance of TrkB^+^ capillary EC following injury, we examined the lungs of young and aged mice during the early resolution phase of bleomycin-induced lung injury (37 days post-bleomycin). As shown in Fig.5J, TrkB^+^ cells were detected in both injured young and aged lungs during this reparative phase, though, these intermediate cells were more abundant in aged lungs in areas exhibiting robust tissue remodeling, suggesting a positive correlation between the number of TrkB^+^ cells and the magnitude of lung injury/fibrosis. Altogether, these *in vivo* data substantiated our transcriptional analysis and highlight TrkB^+^ gCap ECs as a population of alveolar capillary ECs that uniquely emerged in injured lungs and persisted in those of aged mice with sustained fibrosis.

Transient activation of hypoxia-mediated signaling pathways, such as HIF1α, has been shown to contribute to angiogenesis and vascular regeneration in multiple organs (56–60). However, sustained tissue hypoxia due to aberrant vascular remodeling leads to disrupted tissue homeostasis and disrepair (59). To investigate the long-term consequences of the maladaptive capillary EC injury responses with aging, we measured tissue hypoxia in young and aged mouse lungs following bleomycin injury (60 days post-injury) and correlated it with the overall capillary density within each examined area. Pimonidazole was used to detect lung hypoxia as previously described (61). As shown in Supplemental Fig.4, injured aged lungs exhibited elevated hypoxia and reduced number of capillaries compared to injured young lungs, in which the capillary network was largely preserved. These findings suggest that sustained tissue hypoxia in fibrotic aged lungs may be responsible for the persistent YAP/TAZ signaling and the limited gCap EC differentiation following lung injury (Fig.5K).

### Impaired aCap EC differentiation is associated with persistent lung fibrosis in aged mice

aCap ECs are localized in the thin regions of the gas-exchange surface in close contact with ATI cells, through which oxygen diffusion occurs (21). Transcriptionally, aCap ECs express a distinct set of genes encoding for proteins implicated in gas exchange such as carbonic anhydrase 4 (*Car4*), and angiogenesis, such as apelin receptor ligand (*Apln*), kinase insert domain receptor (*Kdr*, encoding for the vascular endothelial growth factor receptor 2, VEGFR2), and endothelin receptor type B (*Ednrb*). Previous studies have shown that gCap ECs differentiate into functional aCap ECs following lung injury (21, 23, 25). Since our data suggested that aged gCap ECs failed to differentiate into mature gCap ECs following lung injury, we wondered whether this abnormality might also affect aCap EC maturation. To assess changes in aCap ECs in response to injury and in aging we sub-clustered this EC subtype and identified three distinct subclusters of aCap ECs including two clusters (2 and 3) present in the lungs of sham animals (young and aged) and one cluster (1) exclusively associated with injured lungs (young and aged) (Fig.6A,B). Notably, this latter cluster encompassed aCap ECs largely derived from fibrotic aged lungs (Fig.6C), suggesting that aCap EC turnover is also compromised with aging.

**Figure 6.**
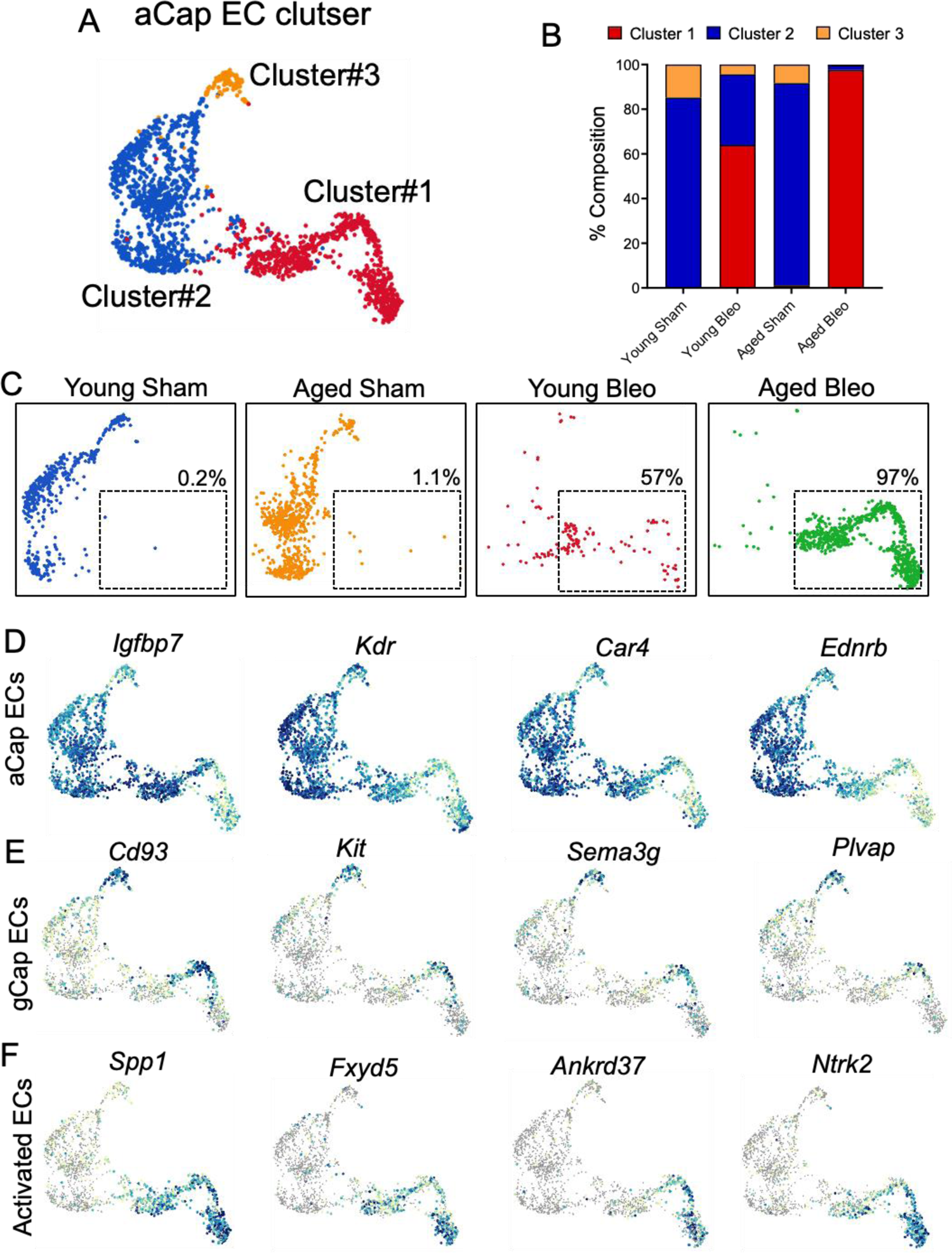
Undifferentiated aCap EC accumulated in fibrotic aged mouse lungs. **(A)** UMAP plot of aCap ECs clusters. **(B)** Composition of aCap EC clusters across different groups. **(C)** UMAP plots displaying aCap EC clusters in young (Blue) and aged (Orange) uninjured lungs, and young (Red) and aged (Green) injured lungs. Intermediate ECs (Cluster #1) co-expressing both aCap and gCap EC marker genes (Red and green) largely accumulate in the lungs of aged mice compared to young ones. **(D)** UMAP plots showing the expression of aCap EC marker genes (Upper), gCap EC marker genes (Middle), and Activated EC marker genes (Bottom) across different clusters.

Gene expression analysis revealed that, though at different levels, all three clusters expressed aCap EC markers, including *Igfbp7*, *Kdr*, and *Car4* (Fig.6D), however, ECs in clusters 1 and 3 co-expressed gCap EC markers (Fig.6E), suggesting that ECs in these two clusters are derived from pre-existing gCap ECs. We also found that, although clusters 1 and 2 shared common gCap EC markers, cluster 1 exclusively expressed markers of cell activation, which are shared with activated gCap EC, including *Spp1*, *Ankrd37*, and *Ntrk2* (Fig.6F), suggesting that ECs in cluster 1 likely derived from aged gCap ECs with compromised differentiation capacity. These findings strongly suggest that, similarly to alveolar epithelial cells, capillary ECs transit into intermediate cell states during lung repair, and aging-associated transcriptional and/or metabolic maladaptation to injury may lead to their commitment failure and aberrant persistence.

### *Ackr1*^+^ venous endothelial cells expand in fibrotic aged mouse lungs

Venous endothelial cells have largely been studied in the context of tissue inflammation and immune cell recruitment both *in vitro* and *in vivo* (*62*). The contribution of venous ECs to lung repair, and more broadly to organ fibrosis, however, is less known. Pulmonary venous ECs are heterogeneous and their properties and functions vary based on their anatomic location (bronchial *vs* pulmonary circulation), and position in the vascular tree (veins *vs* venules) (63–65).

To study the impact of aging and bleomycin-induced lung injury on pulmonary venous EC responses, we sub-clustered venous EC based on the expression of previously identified markers, including *Nrfr2*, *Slc6a2*, and *Amigo2*. Graph-based cluster analysis identified six transcriptional distinct venous EC subtypes (Fig.7A) - clusters 1, 2, and 4 mainly included ECs from uninjured lungs (young and aged), whereas clusters 3, 5, and 6 were predominantly occupied by ECs from injured lungs (young and aged) (Fig.7B-D). While the number of activated venous EC in clusters 3 and 5 was relatively similar between young and aged lungs, with a moderate increase in aged lungs, activated venous ECs in cluster 6 exclusively emerged in aged lungs (Fig.7E). Gene enrichment analysis identified *Atf3*, *Adgrl3,* and *Fabp4*, and as distinctive gene markers of quiescent venous ECs (clusters 1, 2 and 4) whereas *Spp1*, *Ankrd37*, and *Cmah*, were highly expressed in all activated venous ECs (clusters 2, 5, and 6) (Fig.7G). This analysis also uncovered a unique gene expression signature in aged venous ECs from cluster 6, of which *Ackr1* was the most distinctive gene marker (Fig.7H). *Ackr1* encodes for Atypical Chemokine Receptor-1 (ACKR1), a G protein-coupled receptor with a major role in chemokine sequestration, degradation, and transcytosis (66, 67). ACKR1 was also shown to regulate chemokine bioavailability and, consequently, leukocyte recruitment (68, 69). Together with *Ackr1*, we identified other inflammatory-related genes that were highly expressed in venous ECs from cluster 6, including *Selp* and *Tifa* (Fig.7H). *Selp*, encodes for p-selectin, a membrane receptor that mediates the interaction between activated ECs and leukocytes (70), whereas *Tifa* encodes for TRAF Interacting Protein with Forkhead Associated Domain, an adapter protein that plays a key role in the activation of NF-kappa-B signaling pathway (71). We also found that venous ECs in cluster 6 shared numerous inflammatory-related gene markers with those in cluster 4, including *Il1r1*, *Ptgs2, and Serpine1* (Fig.7I), suggesting that this newly emerged venous EC population may be derived from preexisting lung venous ECs with specialized inflammatory functions. This hypothesis was supported by gene ontology analysis on cluster 6 which showed that inflammation and immune cell activation were among the enriched biological processes in this cluster (Fig.7J). In addition, IPA pathway analysis identified several enriched signaling pathways in this venous EC population, including those implicated in vessel remodeling and angiogenesis, such as Rho family GTPase, integrins, VEGF, and axonal guidance (Fig.7K). These findings indicate that a venous EC population exhibiting inflammatory and angiogenic features emerged in fibrotic aged mouse lungs, implicating venous remodeling as a previously unappreciated aging-associated phenomenon with potential repercussions on the development of lung fibrosis.

**Figure 7.**
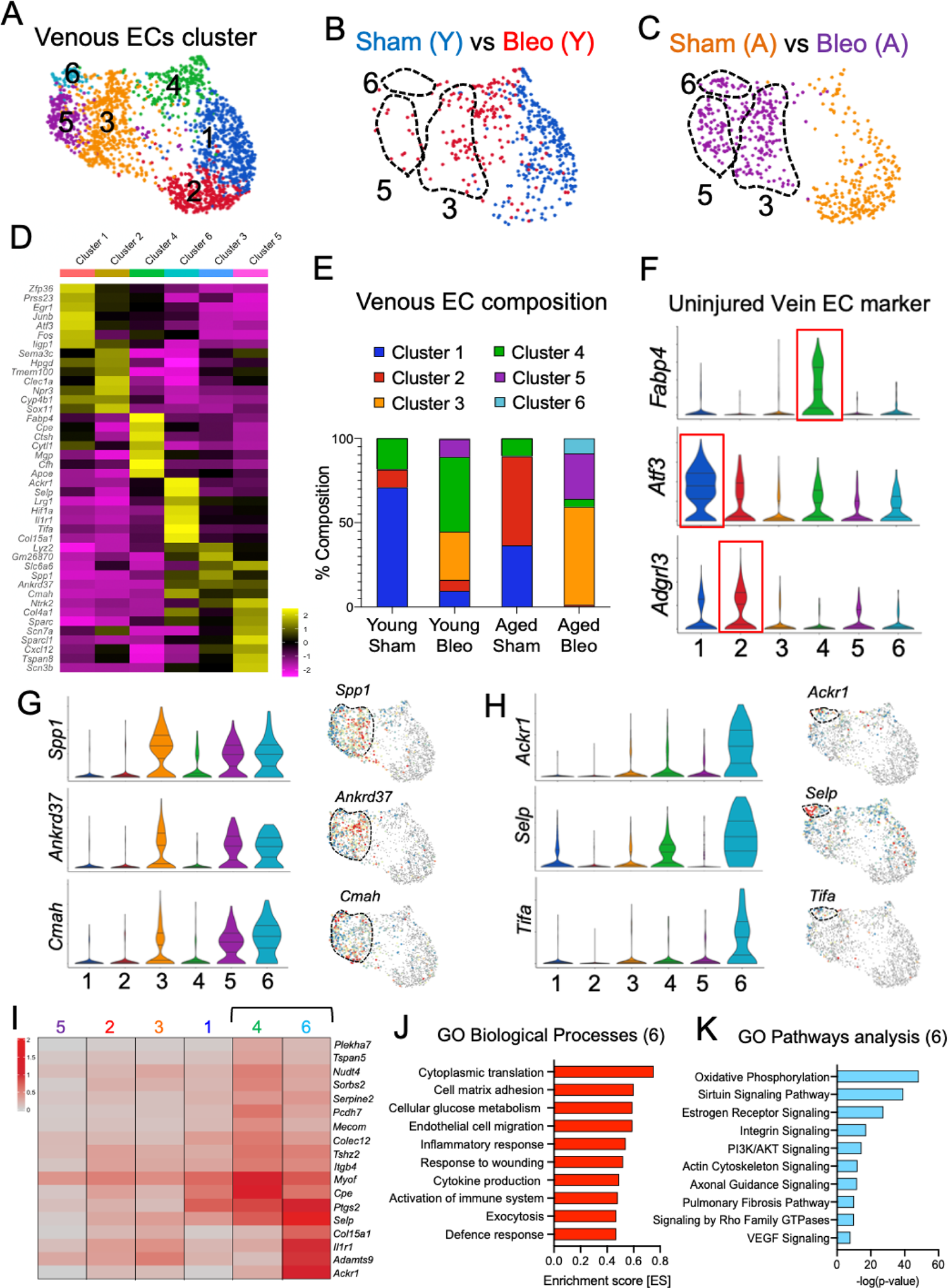
Venous EC remodeling in young and aged mouse lungs following bleomycin challenge. **(A)** UMAP plots showing venous EC clusters. **(B, C)** UMAP plots displaying venous EC clusters in young (Blue) and aged (Orange) uninjured lungs, and young (Red) and aged (Purple) injured lungs. Venous ECs in Clusters 3, 5, and 6 exclusively emerged in bleomycin-injured lungs and their number increased in fibrotic aged lungs compared to young ones. **(D)** Heatmap showing the average expression of venous EC marker genes across different clusters **(E)** Composition of venous EC clusters across different groups. **(F)** Violin plots showing the expression of several marker genes defining different venous EC subtypes at baseline. **(G)** Violin and UMAP plots showing the expression of distinctive marker genes in activated venous ECs (clusters 3, 5 and 6). **(H)** Violin and UMAP plots showing the expression of marker genes defining an activated venous EC population that exclusively emerged in fibrotic aged lungs (clusters 6). **(I)** Heatmap showing the average expression of venous EC markers across different clusters. Genes enriched in cluster #6 largely overlap with those in cluster #4, suggesting that these cell clusters are transcriptionally coupled. **(J, K)** GO analysis shows that genes preferentially expressed in venous ECs from cluster #6 are largely associate with angiogenesis and inflammation.

To shed further light into the pathological venous remodeling associated with lung fibrosis, we immunostained normal and fibrotic lungs form young and aged mice with anti-ACKR1 antibodies (69) and the pan-EC marker CD31. As shown in Fig.8A, in normal mouse lungs, CD31 was strongly expressed in alveolar capillary ECs and weakly expressed in ECs from larger vascular beds. On the contrary, ACKR1 expression was exclusively detected in lung ECs from small venules and large veins. ACKR1^+^ venules were mainly found underneath the epithelial layer of both large bronchi and small bronchiole. Though sporadic, ACKR1^+^ venules were also identified in the alveolar space where they are intimately connected with CD31^+^ capillaries of the pulmonary circulation. Immunostaining analysis of aged mouse lungs with delayed fibrosis resolution showed that ACKR1^+^ venules were closely associated with alveolar regions exhibiting reduced capillary density, alveolar thickening, and immune cell aggregation (Fig.8B). We also found that ACKR1^+^ venules were abundant in bleomycin-treated aged lungs compared to young ones and were closely associated with αSMA^+^ mesenchymal cells (Fig.8C), suggesting that venules-associated stromal cells may be a putative source of myofibroblasts following lung injury.

**Figure 8.**
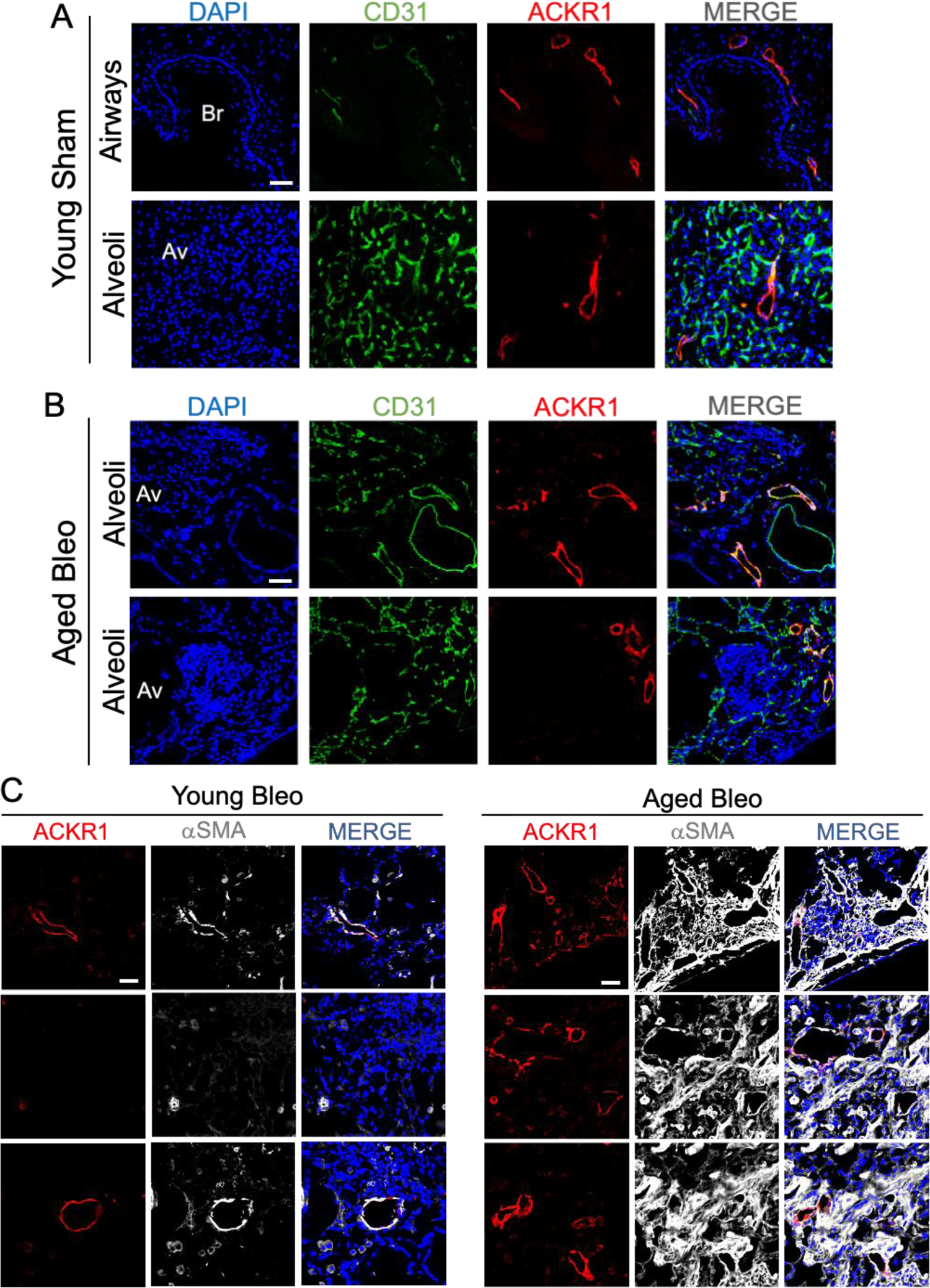
ACKR1^+^ venous ECs reside in bronchial and alveolar regions of the lung and are closely associated with areas exhibiting fibrotic remodeling in aged lungs. **(A)** Immunofluorescence staining showing ACKR1 (venous EC marker) and CD31 (Pan-endothelial marker) expression in lung airways and alveoli at baseline and **(B)** in bleomycin-treated aged lungs. **(C)** Immunofluorescence staining using antibodies against ACKR1 and α-SMA in young and aged lungs following bleomycin injury. ACKR1 positive venous EC accumulates in lung areas exhibiting extensive remodeling. Scale bar 50 µm.

### ACKR1^+^ venous endothelial cells expand in human fibrotic lungs

To translate our observations to human disease, we carried out immunostaining analysis on human lungs isolated from patients with IPF as well as on those form healthy donors. As shown in Fig.9A, in normal human lungs ACKR1^+^ ECs were observed in small venules and veins around bronchi, bronchiole, and in the alveoli, and were focally surrounded by adventitial stromal cells expressing high levels of Collagen-I and αSMA. Intriguingly, Col1^+^ αSMA^+^ stromal progenitor cells were recently identified in the peribronchial and alveolar regions of mouse and human lungs and were shown to give rise to pathogenic myofibroblasts in a mouse model of lung fibrosis (72), further suggesting that ACKR1^+^ EC/stromal cell crosstalk may be implicated in myofibroblasts appearance.

**Figure 9.**
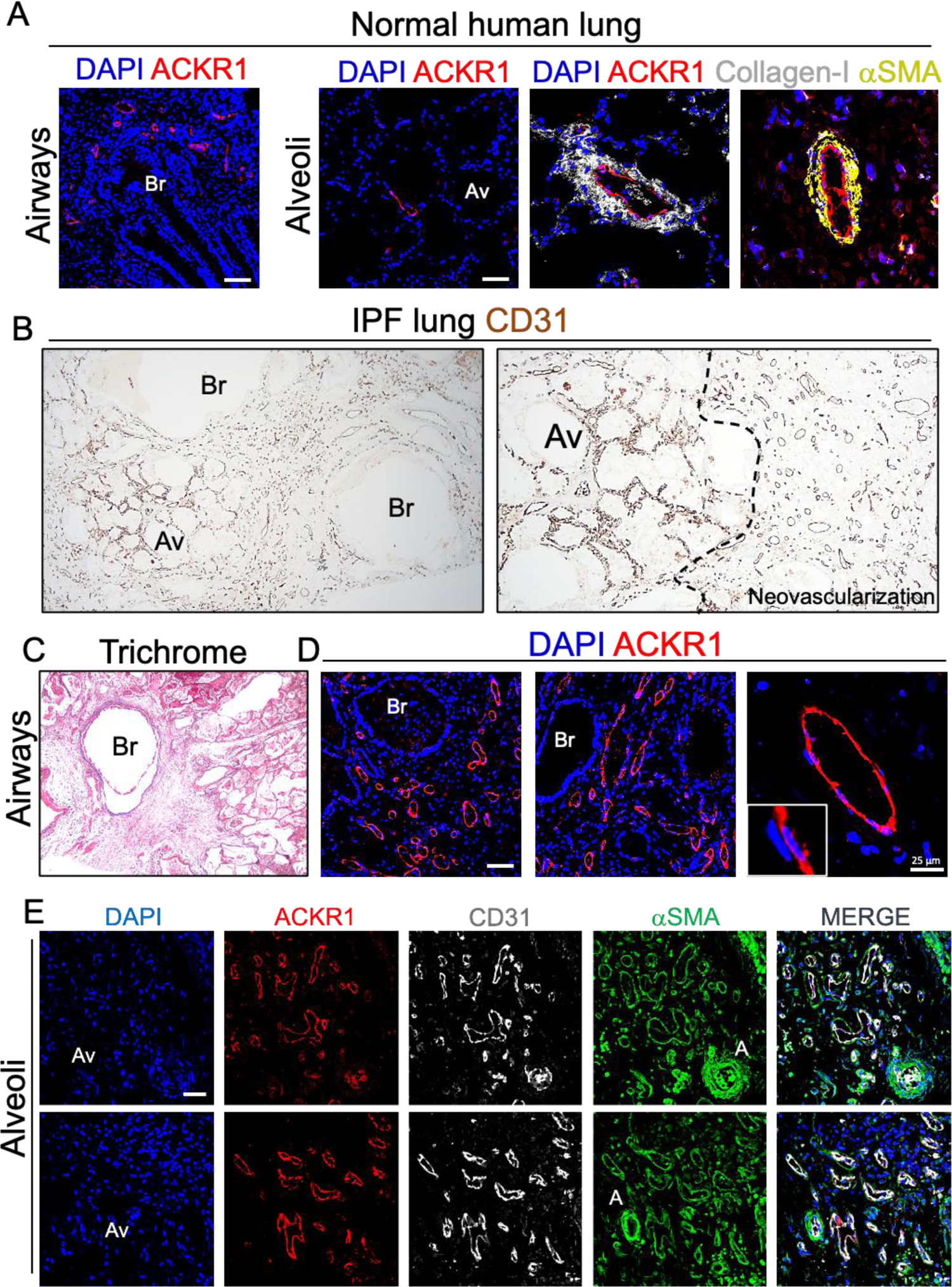
ACKR1^+^ venous ECs expand in human fibrotic lungs. **(A)** Immunofluorescence staining of normal human lungs using antibodies against ACKR1, Collagen-I, and αSMA. ACKR1 is mainly expressed in peribronchial and alveolar venous ECs. Perivascular cells surrounding ACKR1 positive venous ECs strongly expressed Collagen-I and αSMA. **(B)** Immunohistochemistry of human IPF lung sections showing normal-looking alveoli surrounded by highly vascularized (CD31 positive) fibrotic areas. **(C)** Trichrome Masson staining of human IPF lung sections showing peribronchial fibrosis. **(D)** Immunofluorescence staining of human IPF lung sections showing ACKR1^+^ veins that surround fibrotic bronchiole and expand toward the alveolar space **(E)** ACKR1 positive veins are found in the fibrotic alveolar space and surrounded by αSMA^+^ mural cells. Scale bar 50 µm.

The vascular abnormalities described in IPF lungs have generated extensive debates and controversy over the last two decades (73–75). Early studies, in fact, have reported both capillary loss as well as increased vessels density in IPF lungs (11, 73, 76), suggesting that these divergent vascular abnormalities may co-exist in the fibrotic lung, and may reflect a diverse endothelial maladaptation to injury in different lung EC subtypes across different locations. To shed further light into the pathological vascular remodeling and spatial endothelial heterogeneity associated with human lung fibrosis, we first immunostained IPF lungs with antibodies against the pan-endothelial cell marker CD31. As shown in Fig.9B, CD31 staining highlighted endothelial cells across different lung locations, including small alveolar capillaries and large vessels around bronchi. Notably, the alveolar parenchyma in these diseased lungs was largely replaced by vascularized fibrotic tissue, which enveloped terminal bronchi and extended diffusely into the adjacent lung parenchyma resulting in distortion and replacement of alveolar structures (Fig.9B,C). Compared to intact parenchyma, areas of fibrosis contained a reduced number of vessels which were however larger than the capillaries present in the alveolar septa. Immunofluorescence analysis using antibodies against CD31 and ACKR1 revealed that most newly formed vessels within this fibrotic tissue were positive for both the pan-endothelial cell marker CD31 and for the venous cell marker ACKR1 and were therefore of venous origin (Fig.9D). Furthermore, alveolar capillaries (CD31^+^ ACKR1^-^) were largely absent in this fibrotic tissue which was mainly populated by venous ECs and αSMA^+^ stromal cells (Fig.9E), demonstrating that, similarly to fibrotic aged mouse lungs, IPF lungs were characterized by aberrant endothelial cell turnover resulting in ACKR1^+^ venous EC expansion. Intriguingly, ACKR1 marks venous ECs of both bronchial and alveolar circulation (Fig.9A), suggesting that ACKR1^+^ venous ECs in IPF lungs may originate from either vascular bed. Developing novel reporter mouse models to trace different populations of venous ECs in fibrotic lungs is paramount to shed light on the origin as well as the contribution of these pathogenic venous ECs to lung scarring.

Our immunostaining analysis of IPF lungs also revealed the presence of several vascular abnormalities involving venous ECs. For example, we detected ACKR1^+^ venous channels in the fibrotic intima of numerous arteries (Figure Supplement 4). These fibrotic vessels also showed medial thickening and αSMA^+^ cells within the thickened intima. While we cannot determine the endothelial origin of ACKR1^+^ veins and the pathogenic mechanisms associated with their appearance in the arterial wall, we speculate that vasa vasorum, which are small vessels residing in the outer layer of arteries (77), may have penetrated the vessel wall through the media leading to neovascularization of the intimal layer. Notably, proliferation of vasa vasorum has been previously associated with neovascularization of atherosclerotic lesions in arterial vessels (78). Inflammatory and hypoxic stimuli originating from the fibrotic intima may have stimulated angiogenic sprouting of ACKR1^+^ venous ECs from the outer arterial layer into the media. Vascular malformations involving arteries and veins, such as arteriovenous shunts, have been previously described in IPF lungs (79, 80), however, the endothelial origin and the mechanisms implicated in these vascular aberrations have remained largely unexplored. Altogether, our findings demonstrated that vascular endothelial turnover is compromised in IPF lungs favoring the expansion of venous ECs while restricting that of alveolar capillaries. Fibrosis of the lung parenchyma in IPF can be patchy, with fibrotic areas alternating with normal-looking parenchyma (81). To link the accumulation of venous EC to the degree of fibrosis, we evaluated the number of venous ECs and fibroblasts in various IPF lung areas using FACS analysis. Based on gross tissue assessment, we selected three different lung areas with various degrees of fibrosis consolidation (high, medium, and low), and carried out a FACS analysis using antibodies against the pan-endothelial cell marker CD31, the venous markers ACKR1 and p-Selectin, and the mesenchymal cell markers Thy-1 and CTHRC1. Of note, Thy-1 was previously identified as a pan-fibroblast marker in the lung, and CTHRC1 was found to be highly expressed in scar-forming fibroblasts in fibrotic mouse and human lungs (72, 82). First, we found that the number of CD31^+^ cells was reduced in fibrotic areas (4.8%) relative to healthy-looking ones (10%), supporting our previous findings that lung ECs are reduced in IPF lungs (9, 11), and that this abnormality inversely correlated with the magnitude of fibrosis. We also found that although the overall number of CD31^+^ ECs was reduced in fibrotic areas, the number of venous ECs (ACKR1^+^ P-selectin^+^) was relatively higher in these areas compared to less fibrotic ones, and this anomaly was accompanied by an increased number of Thy-1- and CTHRC1-expressing fibroblasts (Fig.10).

**Figure 10.**
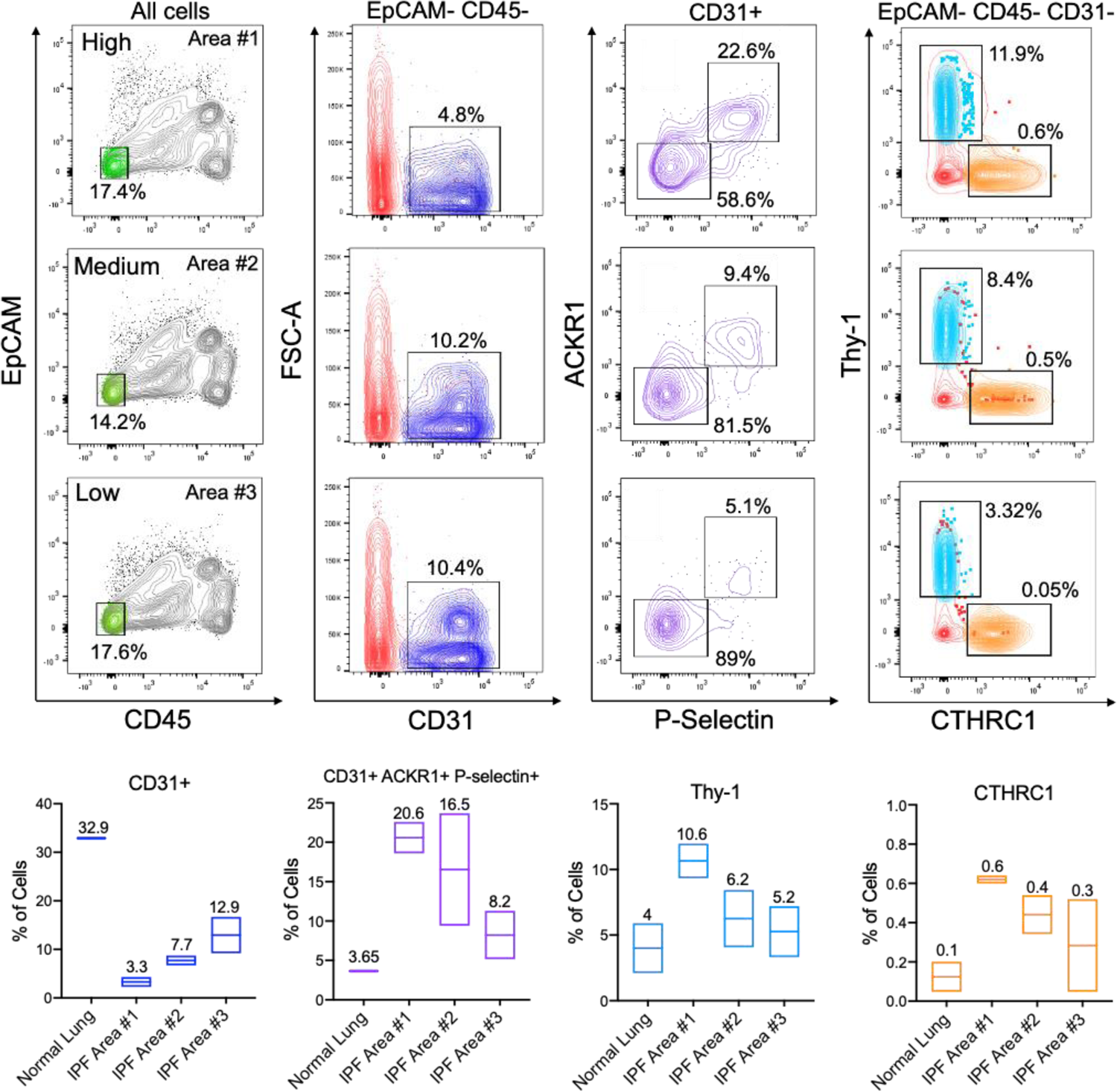
ACKR1^+^ venous ECs expand in lPF lung regions containing elevated number of pathogenic myofibroblasts. FACS analysis of a human IPF lung across different fibrotic regions. Antibodies used for this analysis include those against the epithelial cell marker EpCAM, the immune cell marker CD45, and the endothelial cell marker CD31. Antibodies against ACKR1 and P-Selectin were used to detect venous ECs. ACKR1^+^ P-Selectin^+^ venous ECs were enriched in fibrotic areas containing a high number of pathogenic myofibroblasts (Thy1^+^ CTHRC1^+^).

## Discussion

Lung regeneration is critical to restore organ homeostasis after injury and this function deteriorates with aging (1, 2). Dysfunctional or inadequate regeneration has been associated with disrepair and sustained lung scarring (83, 84), suggesting that loss of regenerative signals with aging contributes to perpetuating fibrogenic signals into the lung resulting in exuberant ECM protein deposition and deterioration of organ functions (1, 20). In this work, we investigated the cellular and transcriptional alterations accompanying persistent lung fibrosis in aged mice following bleomycin injury and compared them to those observed in young mice in which fibrosis typically resolves. Our scRNA-seq analysis revealed numerous aging-associated cellular and transcriptional alterations in response to bleomycin injury that affected epithelial and endothelial cell lineages, including the appearance of activated intermediate states that persisted in fibrotic aged and IPF lungs. Given that interstitial fibroblasts are intimately connected with both epithelial and endothelial cells in the alveolus, loss or limited alveolar regeneration with aging due to dysfunctional differentiation may directly impair fibroblast behavior and contribute to chronic scarring.

Our scRNA-seq analysis gave us the opportunity to investigate cellular alterations associated with the persistent lung fibrosis of aged mice. Our observations suggest that *Pdgfra*^+^ alveolar fibroblasts are the major source of activated fibroblasts in response to bleomycin injury in both young and aged lungs, exhibiting high expression of genes typically associated with tissue scarring, including collagen and matrix remodeling genes. These observations corroborated previous studies in mouse and humans showing PDGFRα^+^ cells as alveolar progenitor cells capable of sensing and responding to local injury by producing large amount of matrix proteins which often persist in chronic settings (85–87). While we did not observe any obvious differences in the number of activated alveolar fibroblasts between young and aged lungs at our selected timepoint (30 days post bleomycin), those derived from aged lungs exhibited the highest expression of genes implicated in fibrosis development, including ECM- and glycolysis-related genes. These analyses not only supported previous observations implicating alveolar fibroblasts as the major source of collagen-producing cells post lung injury, but also substantiated our bulk RNA-seq analysis (10) demonstrating greater fibroblast activation with aging.

During lung fibrosis multiple aberrant signaling pathways in parenchymal cells have been proposed to contribute to a complex cellular interplay resulting in persistent fibroblast activation, including cellular senescence, oxidative stress, and mitochondrial dysfunction (1, 7, 17, 19, 20). A failure to proper differentiate has also been proposed as a putative parenchymal aberration leading to the sustained fibroblast activation with aging (1, 19, 88). For example, stress from genetic or environmental factors can compromise the ability of ATII cells to differentiate into ATI cells, resulting in defective epithelial maintenance (27, 89). Studies have suggested that ATII cells transit into a transcriptionally-defined intermediate state preceding the regeneration of ATI cells following lung injury (27). These intermediate cells, which are marked by high expression of Krt8^+^, exhibited transcriptional features of cellular senescence and persisted in human lungs with fibrosis (27). Our single cell transcriptomic analysis corroborated these previous findings and identified intermediate Krt8^+^ epithelial cells that co-expressed both ATII and ATI markers.

Intriguingly, we discovered that within this intermediate cell population, the number of cells expressing ATI markers was reduced in aged lungs compared to young ones, in which the expression of ATI markers prevailed, suggesting that ATII cell differentiation is compromised with aging. The reduced YAP/TAZ gene expression observed in aged intermediate cells also suggest that the Hippo signaling pathway, which plays critical functions in ATII differentiation (34, 36, 90), is aberrantly regulated with aging. Given the association between YAP/TAZ and mechanotransduction, our observations raise interesting questions about how changes in lung mechanics that associate with age (91) may contribute to aberrant transitional states that persist with injury.

Similar to ATII cells, gCap ECs act as progenitor cells that can differentiate into aCap ECs in response to lung injury (21, 23, 25). Interestingly, our data also support a role for YAP/TAZ in gCap EC differentiation, suggesting that the Hippo signaling may regulate alveolar morphogenesis in multiple cell types in a coordinated fashion. Our results also showed a strong association between the YAP/TAZ transcriptional program and gCap EC responses to injury. Pharmacological activation of YAP/TAZ in capillary ECs *in vitro* promoted the expression of markers associated with endothelial de-differentiation, resembling the intermediate state observed in activated gCap EC *in vivo*. Intriguingly, previous studies have shown that activation of YAP/TAZ signaling in various cell types, including those in the lung epithelium, promotes stem cell activity and prevents differentiation (34, 90, 92, 93).

Our finding together with the existing literature support the premise that YAP/TAZ direct cell fate transitions in both the lung epithelium and endothelium, and aberrant regulation of these processes contribute to the accumulation of intermediate undifferentiated cells in fibrotic aged lungs. Notably, our data suggest that persistent YAP/TAZ promote gCap EC activation with aging leading to aberrant gCap EC differentiation. Age-related increases in stiffness have been reported in parenchymal and vessel compartments of the lung (91), leading us to speculate YAP/TAZ activation in response to mechanisms that control stiffness homeostasis results in sustained activation of this pathway. Given that intermediate gCap ECs exhibited increased expression of numerous HIF1α target genes and fibrotic aged lungs exhibited elevated hypoxia, another related possibility is that hypoxia-associated signaling pathways may alter YAP/TAZ responses to drive intermediate gCap ECs accumulation with aging.

Our transcriptional analysis together with lineage tracing identified TrkB as a YAP/TAZ-regulated gene that was distinctly expressed by intermediate activated gCap ECs. Previous studies showed that TrkB together with its ligand brain-derived neurotrophic factor (BDNF), orchestrate neural cell activation and survival (94, 95). Previous studies demonstrated that BDNF/TrkB signaling is critical during developmental angiogenesis and vessels maturation (96–98), and activation of this pathway was shown to support angiogenesis by influencing vascular endothelial growth factor receptor-2 (VEGFR2) expression (99). These findings suggest that TrkB together with specific endothelial signaling pathways, such as VEGF pathway, may be critical for the differentiation and maturation of gCap ECs. Interestingly, the TrkB ligand BDNF is strongly expressed by intermediate ATII cells suggesting functional crosstalk between different intermediate cell populations in supporting alveolar regeneration following injury. Given that our scRNA-seq analysis revealed abnormal ATII transcriptional responses to injury, including reduced expression of *Bdnf* and *Vegf* (data not shown), and accumulation of undifferentiated gCap ECs with aging, these findings suggest that aging-mediated aberrant epithelial cell responses to injury may negatively impact gCap EC differentiation. In support of the dysfunctional gCap EC differentiation with aging, our FACS analysis showed that the number of capillary ECs expressing the gCap marker TEK was strongly reduced in IPF lungs compared to healthy lungs, further implicating the loss of gCap EC identity as a pathogenic feature of fibrotic lungs with compromised regenerative capacity. Restoring regenerative signaling pathways in lung progenitors, including ATII and gCap ECs, may provide avenues for enhancing alveolar repair in elderly suffering from chronic lung disorders, including IPF.

Our single cell transcriptomic analysis also revealed the appearance of multiple populations of ECs derived from venous EC lineages in both injured young and aged lungs. To our knowledge, no studies in mice have previously investigated the distribution, localization, as well as the contribution of venous EC to lung injury responses. Our clustering analysis of lung venous ECs led to the identification of three distinctive venous EC populations in uninjured mouse lungs, and each of them was characterized by the expression of distinct marker genes, suggesting that lung venous ECs may serve different functions in different lung locations and vascular districts where they are exposed to regionally distinct differentiation signals. The gene expression signatures of venous ECs were profoundly altered following injury in both young and aged lungs as demonstrated by the acquisition of injury markers that were not expressed by quiescent venous ECs. These distinctly expressed marker genes, including *Spp1*, *Ankrd37*, and *Ntrk2,* were also expressed by activated gCap ECs, suggesting that both endothelial lineages converge on common transcriptional states before returning to quiescence.

Our analysis revealed that the composition of venous ECs that emerge with injury are different between young and aged lungs, with aged lungs exhibiting a distinct venous EC population characterized by the expression of the marker gene *Ackr1* and the upregulated expression of genes implicated in inflammation, suggesting that these cells may orchestrate inflammatory responses that contribute to the lung fibrotic milieu. Consistent with this inflammatory function, venous remodeling was mainly observed in dense fibrotic regions of aged mouse lungs, while this feature was less pronounced in injured young mice. Similarly, in IPF lungs ACKR1^+^ veins were also abundant in fibrotic areas and largely surrounded by αSMA^+^ cells, suggesting a pathogenic function for this venous EC population. This hypothesis was reinforced by our FACS analysis in IPF lungs demonstrating a positive correlation between the number of myofibroblasts and that of venous ECs in lung areas with different degree of fibrosis. Recently, a population of venous ECs marked as *COL15a1*^+^, and whose location in healthy human lungs was restricted to bronchi and bronchioles (65), was found in distal alveolar regions of IPF lungs (100), suggesting that venous EC-mediated neovascularization may contribute to disease progression. Intriguingly, in our mouse study we found that *Col15a1* was also enriched in *Ackr1^+^* venous ECs, however, the expression of this gene was not as distinctive as *Ackr1* and it was also detected in other activated venous ECs. Given that ACKR1^+^ venules were found to reside in different lung locations and circulatory systems (bronchial and pulmonary circulations) (65), we cannot conclusively determine from our study which venous EC population is involved in the aberrant vascular remodeling associated with persistent fibrosis in aged lungs. Altogether, these findings shed new light into the putative role of venous remodeling on bleomycin-induced lung fibrosis, and open new research opportunities to explore how lung injury and aging influence endothelial cell behavior in different vascular beds.

## Material and methods

### Animals

Male young (2 months old) and aged (18 months old) C57BL6 mice were purchased from Jackson Laboratory (Bar Harbor, ME). All mice had access to food and water ad libitum and were on a 12 h/12 h light/dark cycle, ambient temperature 77–78 °F and humidity 46–49%.

### Cell culture and treatments

Human lung microvascular endothelial cells (HLMECs) were purchased from Cell Applications (San Diego, CA, USA) or ANGIO-PROTEOMIE (Worcester, MA, USA) and maintained in endothelial cell growth basal medium supplemented with microvascular endothelial cell growth kit. Cells were treated with 2 µM of TRULI (Selleckchem), a LATS inhibitor and YAP/TAZ signaling activator, for 48 hrs. All the experiments were performed with cells within 3-5 passage.

### Mouse model of bleomycin-induced lung injury

All animal experiments were carried out under protocols approved by the Boston University Institutional Animal Care and Use Committee (IACUC) and conformed to the ARRIVE guidelines. Bleomycin was delivered to the lungs as previously described (11, 101). Mice were anesthetized with ketamine/xylazine solution (100 mg/kg, Cat# 0143-9509 and 10 mg/kg, Cat# 59399-110 respectively), and injected intraperitoneally. About 1 U/Kg bleomycin (APP Pharmaceutical, LCC Schaumburg, IL, USA) or PBS (50ul) were intratracheally delivered using an insulin syringe (). Body weight was monitored daily.

### Lineage tracing experiments

The following mouse lines were used: Aplnr-CreER (Tg(Aplnr-cre/ ERT2)#Krh) (21) (Kindly provided by Dr. Kristy Red-Horse, Stanford University). Rosa26-mtdTomato/mGFP mice (B6.129(Cg)-Gt(ROSA)26Sortm4(ACTB-tdTomato,-EGFP)Luo/J (Jax #007676). Lineage-tracing was induced by tamoxifen injection (5 injections of 75mg tamoxifen/kg body weight, daily).

### Human lung tissues

Lung tissues from patients with IPF were obtained from explanted lungs obtained at the time of transplantation (all with informed consent and approved by the University of Michigan, Ann Harbor, MI) and from an autopsy (VA Medical Center, Seattle, WA). Diagnoses of patients with IPF were established by clinical pathological criteria and confirmed by multidisciplinary consensus conference. Normal control lungs were obtained from deceased donors (Gift of Life, Michigan) whose lungs were deemed unsuitable for transplant.

### Hypoxia detection by Pimonidazole

Hypoxyprobe™-1 RED PE (Cat# HP-1 RED PE Mab-1, Hypoxyprobe, Inc; Burlington, MA, USA) was used to characterize hypoxia within the lungs of young and aged mice either sham or injured with bleomycin. 60 days after administration of bleomycin, mice were injected with pimonidazole (60 mg/kg) intraperitoneally 60 min before harvesting of the lungs. For the harvest, mice were anaesthetized with ketamine/xylazine solution, perfused via left ventricle with cold PBS and inflated with a solution of OCT and PBS (50%/50%), followed by embedding of the samples in OCT compound (Tissue-Tek 4583; Sakura Finetek Japan, Co. Ltd, Tokyo, Japan). Tissue sections (5 mm) from each block were cut in a cryostat at −21 C and mounted onto Vectabond-coated slides (Vector Laboratories, Peterborough, UK), fixed in cold acetone for 10 minutes and incubated with an anti PECAM-1 rat primary antibody (BD Biosciences, San Jose, CA, USA, Cat# 550274) (1:100 in PBS with 1% BSA and 5% goat serum) overnight at 4 degrees. Sections were then incubated for 1 hour with a fluorescence-conjugated secondary antibody Alexa Fluor 647 (1:1000 Thermo Fisher Scientific, Waltham, MA, USA), a secondary anti-pimonidazole antibody HP RED PE (1:120 in PBS with 0.1% BSA) and DAPI to counterstain nuclei. Controls were done by omitting the primary antibodies. Two to 6 regions per mouse were analyzed and mean ± SD was determined.

### Single-cell RNA sequencing and data analysis

Young lungs (1 Sham and 1 Bleo-treated) and aged lungs (2 Sham and 2 Bleo-treated) were harvested and disassociated into a single-cell suspension as previously described (9). Thirty days pos-bleomycin treatment, mice were anesthetized with ketamine/xylazine solution and perfused via the left ventricle with cold PBS 30. The lungs were immediately harvested and minced with a razor blade in a 100 mm petri dish in a cold DMEM medium containing 0.2 mg/ml Liberase DL and 100 U/ml DNase I (Roche, Indianapolis, IN, USA). The mixture was transferred into 15 ml tubes and incubated at 37 °C for 35 min in a water bath under continuous rotation to allow enzymatic digestion. Digestion was inactivated with a DMEM medium containing 10% fetal bovine serum, the cell suspension was passed through a 40 µm cell strainer (Fisher, Waltham, MA, USA) to remove debris. Cells were then centrifuged (500×g, 10 min, 4 °C), and suspended in 3 ml red blood cell lysis buffer (Biolegend, San Diego, CA, USA) for 90 s to remove the remaining red blood cells and diluted in 9 mL PBS after incubation. Cells were then centrifuged (500×g, 10 min, 4 °C) and resuspended in 0.2 ml of 1X PBS buffer containing 0.04% BSA. Cell suspensions were submitted to the Single-Cell Sequencing Core (Boston University) for processing. Cell viability and counts were determined using Countess II Automated Cell Counter. Single cells, reagents, and a single Gel Bead containing barcoded oligonucleotides are encapsulated into nanoliter-sized Gel Bead-inEmulsion using the 10x Genomics GemCode platform (10X Genomics, Pleasanton, CA, USA). Lysis and barcoded reverse transcription of RNAs from single cells is performed as described by 10x genomics. Enzyme fragmentation, A tailing, adapter ligation, and PCR are performed to obtain final libraries containing P5 and P7 primers used in Illumina bridge amplification. Size distribution and molarity of resulting cDNA libraries were assessed via Bioanalyzer High Sensitivity DNA Assay (Agilent Technologies, USA). All cDNA libraries were sequenced on an Illumina NextSeq 500 instrument according to Illumina and 10X Genomics guidelines with 1.4–1.8 pM input and 1% PhiX control library spike-in (Illumina, USA). Sequencing data were processed using 10X Genomics’ Cell Ranger pipeline to generate feature/barcode matrices from raw count data.

Feature and barcode matrices of the samples were imported into BioTuring Browser 3 (Bioturing, San Diego, USA) for analysis. After quality filtering, we obtained 52542 cell profiles, 11767 from young sham, 5893 from young Bleo, 16507 from aged sham and 18375 from aged bleo mice. We perform t-Distributed Stochastic Neighbor Embedding (t-SNE) dimensionality reduction with canonical correlation analysis (CCA) subspace alignment and performed unsupervised graph-based clustering. Analysis of representative marker genes identified clusters of endothelial, epithelial, and mesenchymal cells as well as hematopoietic cells. Indicated cells were selected and further re-clustered for analysis. The criteria for selection of significant differentially expressed genes were: log2 fold change ≤-0.1 or ≥0.1 and Benjamini– Hochberg adjusted p value or FDR ≤0.05. This list of differentially expressed genes was used for investigating enriched canonical pathways and upstream regulators using Core analysis from the Ingenuity Pathway analysis (IPA, Ingenuity® Systems, www.ingenuity.com). P-values were generated in IPA using Fisher’s test (Log2 fold change ≤-0.1 or ≥0.1, p value ≤0.05). p value and activation z score were used for plotting noteworthy canonical pathways and activated upstream regulators respectively.

### Quantitative Real-time PCR

Total mRNA was isolated using Quick-RNA^TM^ Miniprep (Zymo Research, Irvine, CA, USA) followed by Nanodrop concentration and purity analysis. cDNA was synthesized using High-capacity cDNA Reverse Transcription Kit (Applied Biosystems); RT-PCR was performed using PowerUp™ SYBR™ Green Master Mix (Applied Biosystems) and analyzed using a Step-One-Plus Real-Time PCR system (Applied Biosystems).

### RNA interference

Transient RNA interference was performed with siGENOME non-targeting Control siRNA Pool #1 (D-001206-13-05, 25 nM) or Human YAP/TAZ-siRNA (UGUGGAUGAGAUGGAUACA) (102) by using Lipofectamine RNAiMAX reagent (13778075, Thermo Fisher Scientific, Waltham, MA, USA). Cells were harvested after 72 h.

### Immunohistochemistry

Formalin-fixed paraffin-embedded (FFPE) mouse and human lungs were cut in serial sections (7 μm). The FFPE sections were deparaffinized using a standard protocol of xylene and alcohol gradients. Sections were then blocked first with BLOXALL endogenous peroxide blocker (SP-6000-100, Vector Laboratories, Peterborough, UK) and then with 5% goat serum and 2% BSA (Sigma-Aldrich, St. Louis, MA, USA). Staining was performed using the VECTASTAIN Elite ABC HRP kit (PK-6200, Vector Laboratories, Peterborough, UK), the anti-mouse CD31 primary antibody (550274, clone MEC 13.3, BD Biosciences, San Jose, CA, USA, 1:200 dilution), the anti-human CD31 primary antibody (clone JC70, Cell Marque, Millipore Sigma, USA), and the detection with impact DAB (Vector Laboratories, Peterborough, UK). Slides were then dehydrated using a standard protocol and mounted on a coverslip using DPX mountant (Sigma-Aldrich, St. Louis, MA, USA). Masson’s trichrome staining was performed by using a commercially available stain kit (HT15, Sigma-Aldrich, St. Louis, MA, USA).

### Immunofluorescence staining

Human or mouse lung tissue slides (7 μm) were fixed in 3.7% formalin (Sigma-Aldrich, St. Louis, MA, USA), permeabilized in 0.1% Triton X-100 (Sigma-Aldrich, St. Louis, MA, USA), blocked with 5% BSA for 1 h. Mouse lung tissue sections were stained with anti-CD31 antibody (77699, Cell Signaling Technology, Danvers, MA, USA 1:200 dilution), anti-TrkB antibody (AF1494, Novus, Centennial, CO, USA, 1:200 dilution), anti-αSMA antibody (F3777, clone 1A4, Sigma-Aldrich, St. Louis, MA, USA, 1:200 dilution), and anti-mouse ACKR1 (1:200) (kindly provided by Dr. von Andrian). Human lung tissue sections were stained with anti-CD31 antibody (3528, Cell Signaling Technology, Danvers, MA, USA 1:200 dilution), anti-ACKR1 antibody (NB100-2421, Novus, Centennial, CO, USA, 1:200 dilution), anti-Col1α1 antibody (72026, Cell Signaling Technology, Danvers, MA, USA 1:200 dilution) and anti-αSMA antibody (F3777, clone 1A4, Sigma-Aldrich, St. Louis, MA, USA, 1:200 dilution). Cell Sections were stained with fluorescence-conjugated secondary antibodies (Alexa Fluor 488-, Alexa Fluor 555- or Alexa Fluor 647-conjugated, Thermo Fisher Scientific, Waltham, MA, USA, 1:1000 dilution) and DAPI (62248, Thermo Fisher Scientific, Waltham, MA, USA, 1:1000 dilution) to counterstain nuclei.

### FACS analysis

Human normal and IPF lungs were minced with a razor blade in a 100 mm petri dish in a cold DMEM medium containing 0.2 mg/ml Liberase DL and 100 U/ml DNase I (Roche, Indianapolis, IN, USA). The mixture was transferred into 15 ml tubes and incubated at 37 °C for 35 min in a water bath under continuous rotation to allow enzymatic digestion. Digestion was inactivated with a DMEM medium containing 10% fetal bovine serum, the cell suspension was passed through a 40 µm cell strainer (Fisher, Waltham, MA, USA) to remove debris. Cells were then centrifuged (500×g, 10 min, 4 °C), and resuspended in 3 ml red blood cell lysis buffer (Biolegend, San Diego, CA, USA) for 90 s to remove the remaining red blood cells and diluted in 9 mL PBS after incubation. Cells were then centrifuged (500×g, 10 min, 4 °C) and resuspended in 0.2 ml of FACS buffer (0.05% BSA, 0.5 mM EDTA pH 7.4 in PBS). Single cell suspension was stained with antiCD45:Pacific blue (368539, clone 2D1, Biolegend, San Diego, CA, USA, 1:200 dilution), anti-EpCAM:BV650 (324225, colon 9C4, Biolegend, San Diego, CA, USA, 1:200 dilution), anti-CD31:APC/Cyanine7 (303119, clone WM59, Biolegend, San Diego, CA, USA, 1:200 dilution), anti-Tek:PE (334205, clone 33.1, Biolegend, San Diego, CA, USA, 1:100 dilution), anti-ACKR1 (NB100-2421, Novus, Centennial, CO, USA, 1:50 dilution), anti-P-Selectin (NB100-65392, Novus, Centennial, CO, USA, 1:50 dilution), anti-CTHRC1 (PA5-49638, Invitrogen, Waltham, MA, USA, 1:50 dilution), anti-rabbit IgG:FITC (406403, clone Poly4064, Biolegend, San Diego, CA, USA, 1:200 dilution), anti-goat IgG:PE (405307, clone Poly4053, Biolegend, San Diego, CA, USA, 1:200 dilution) and DAPI (D3571, Thermo Fisher Scientific, Waltham, MA, USA, 1:10,000 dilution) for 15 minutes on ice. Cells were washed with ice-cold FACS buffer twice and analyzed by BD LSR II (BD Biosciences, San Jose, CA, USA). FACS analysis was performed with the following strategy: debris exclusion (FSC-A by SSC-A), doublet exclusion (SSC-W by SSC-H and FSC-W by FSC-H), and dead cell exclusion (DAPI by FSC-A). Data were analyzed with FlowJo version 10.8.0 software (Tree Star Inc., Ashland, OR, USA).

### Statistical analysis

Individual data points are shown in all plots and represent data from independent mice, cells, or biological replicates from cell culture experiments. Variables are summarized as mean and SD, with a statistical comparison between two groups performed using Student’s t-test. All analyses and plots were generated using GraphPad Prism 9.4.1 (La Jolla, CA, USA) with statistical significance defined as p < 0.05.

### Data availability

scRNA-seq data generated in this study will be available through the Gene Expression Omnibus (GEO) under accession numbers that will be provided upon acceptance of the manuscript.

## Acknowledgements

We thank Tianmu Hu and Yuriy Alekseyev (Boston University) for supporting bioinformatics analysis. We thank Dr. David Rogers, Pathology and Laboratory Medicine, VA Puget Sound Health Care System, Seattle, WA for his help in providing histologic sections of postmortem lung tissue. We also thank Dr. Kristy Red-Horse (Department of Biology, Stanford University) for sharing the Aplnr-CreER(T) mouse line. We gratefully acknowledge support of this work by the National Institutes of Health (NIH) grants R01HL142596 (G.L.), R01HL158733 (G.L.), R01HL124392 (X.V.), T32HL007035 (T. X. P.) and support from The Evans Center for Interdisciplinary Biomedical Research ARC on “Connecting Tissues and Investigators, Fibrosis in Pathology” at Boston University.

## Author contributions

G.L. and X.V., conceptualized the study and acquired funding. T.X.P., A.R., J.L., J.H., J.S., K.N., N.C., performed the experiments. T.X.P., G.L., A.R., X.V., T.D.M., analyzed and visualized the data. A.T. and U.H.V.A. developed the antibody against mouse ACKR1. The manuscript was drafted by G.L. and A.R., and revised by X.V., M.T., R.F.N., A.M.B., S.K.H., N.C., U.H.V.A. Human lung samples were procured by S.K.H. and R.F.N. All Authors participated in manuscript preparation and provided final approval of the submitted work.

---

**Supplement Figure 1.**
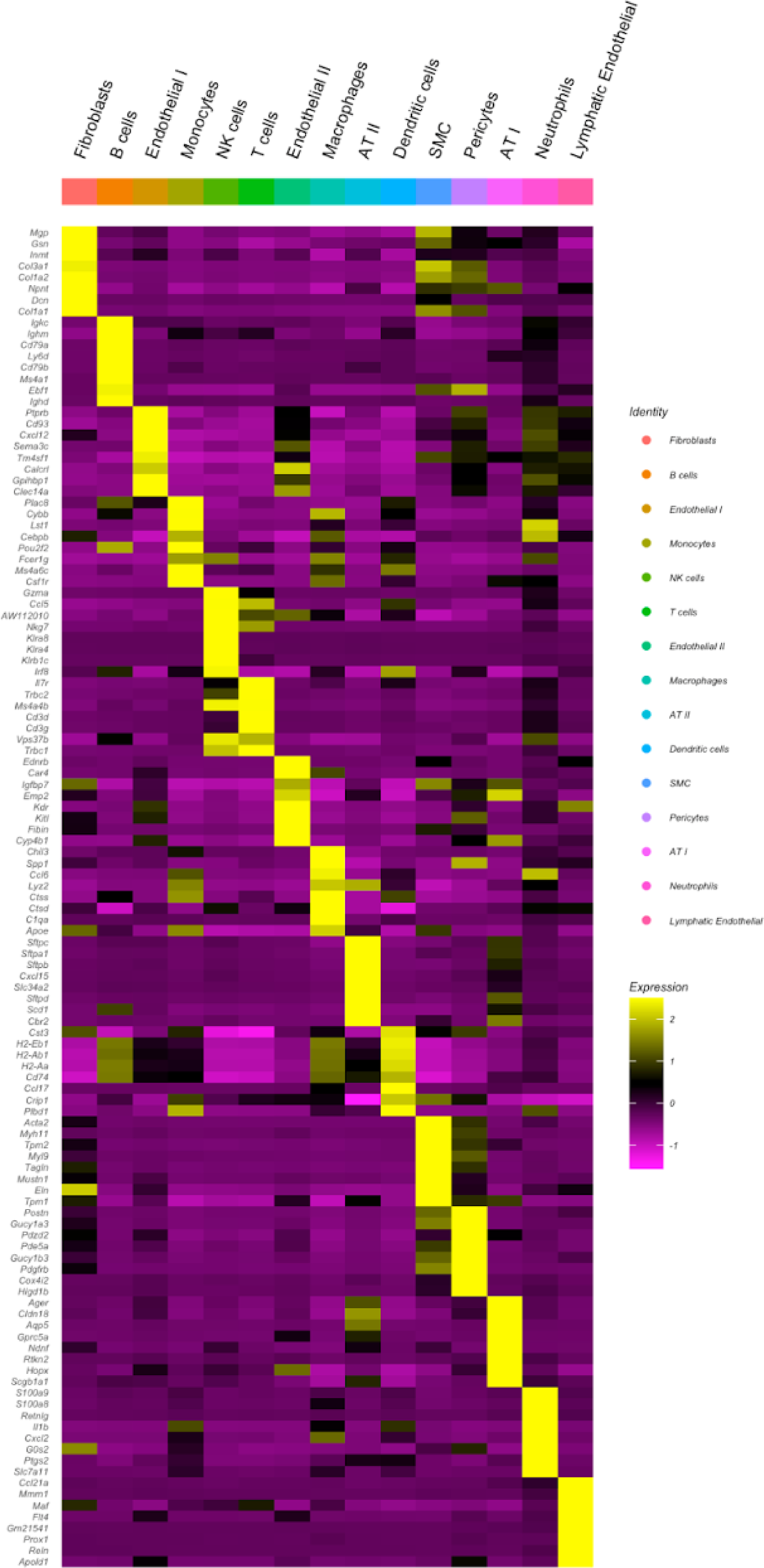
Heat map of marker genes for all identified cell types. Each cell type is defined by the top eight most expressed genes. Each column represents the average expression value for each lung sample, hierarchically grouped by cell type.

**Supplement Figure 2.**
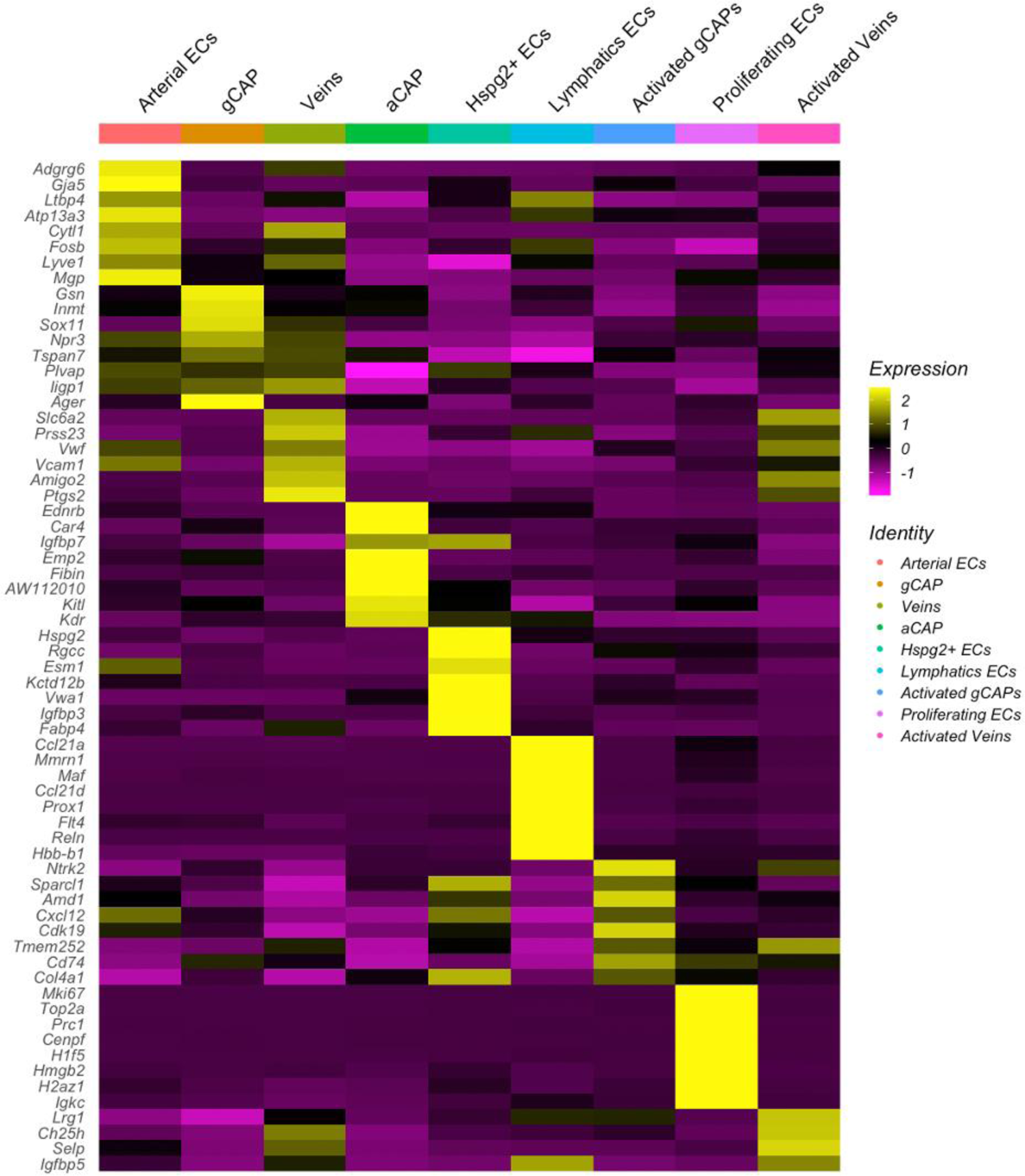
Heat map of marker genes for all endothelial cell types Each EC lineage is defined by the top eight most expressed genes. Each column represents the average expression value for each lung sample, hierarchically grouped by EC type.

**Supplement Figure 3.**
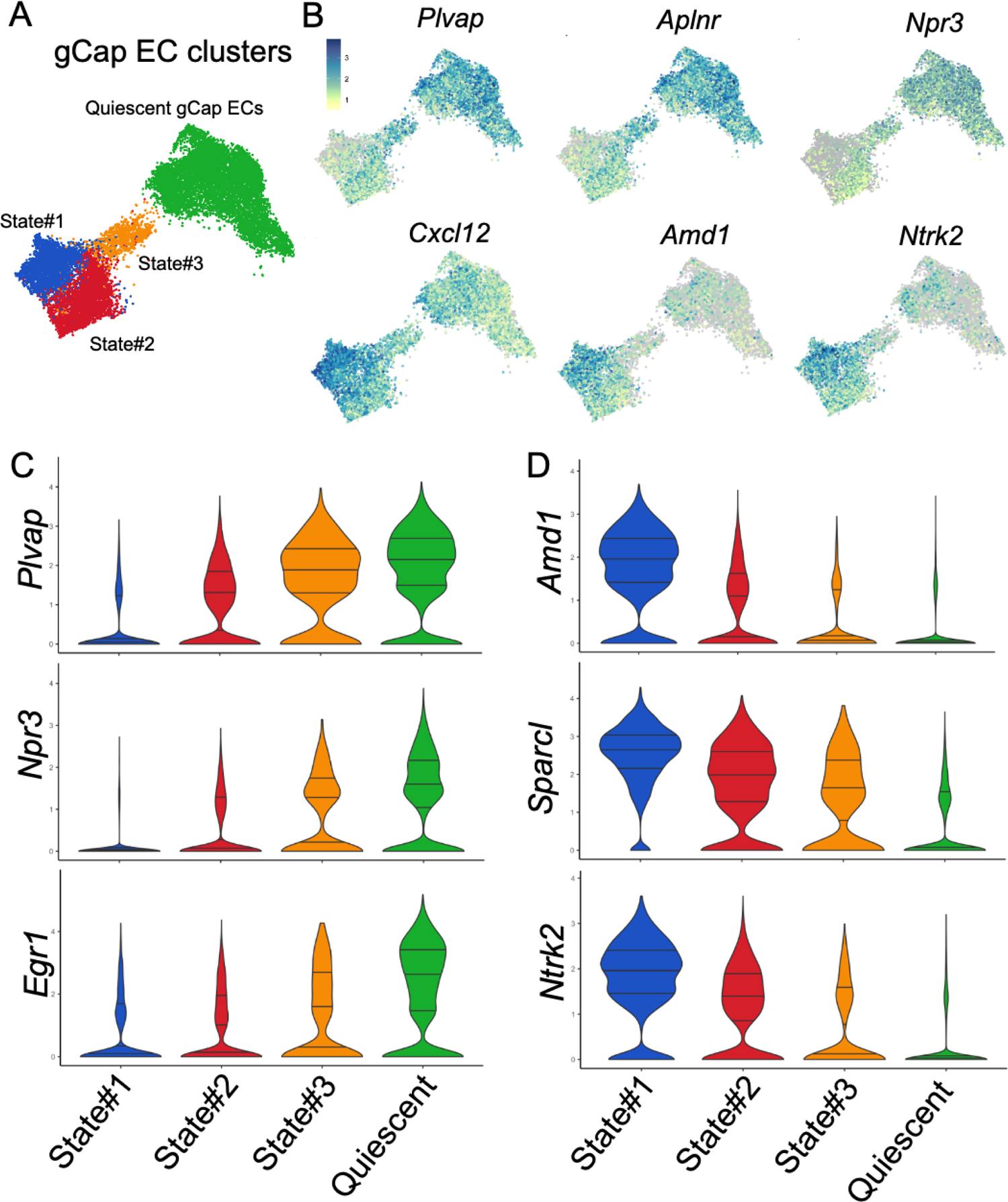
Activated gCap EC clusters are transcriptionally coupled and represent distinct but potentially connected intermediate states. (A) UMAP plots showing a quiescent gCap EC cluster (Green) and three clusters representing distinct transcriptional states of activated gCap EC (B) UMAP plots showing the expression of gCap EC marker genes across different clusters (C) Violin plots of the expression of marker genes in different gCAP EC differentiating states.

**Supplement Figure 4.**
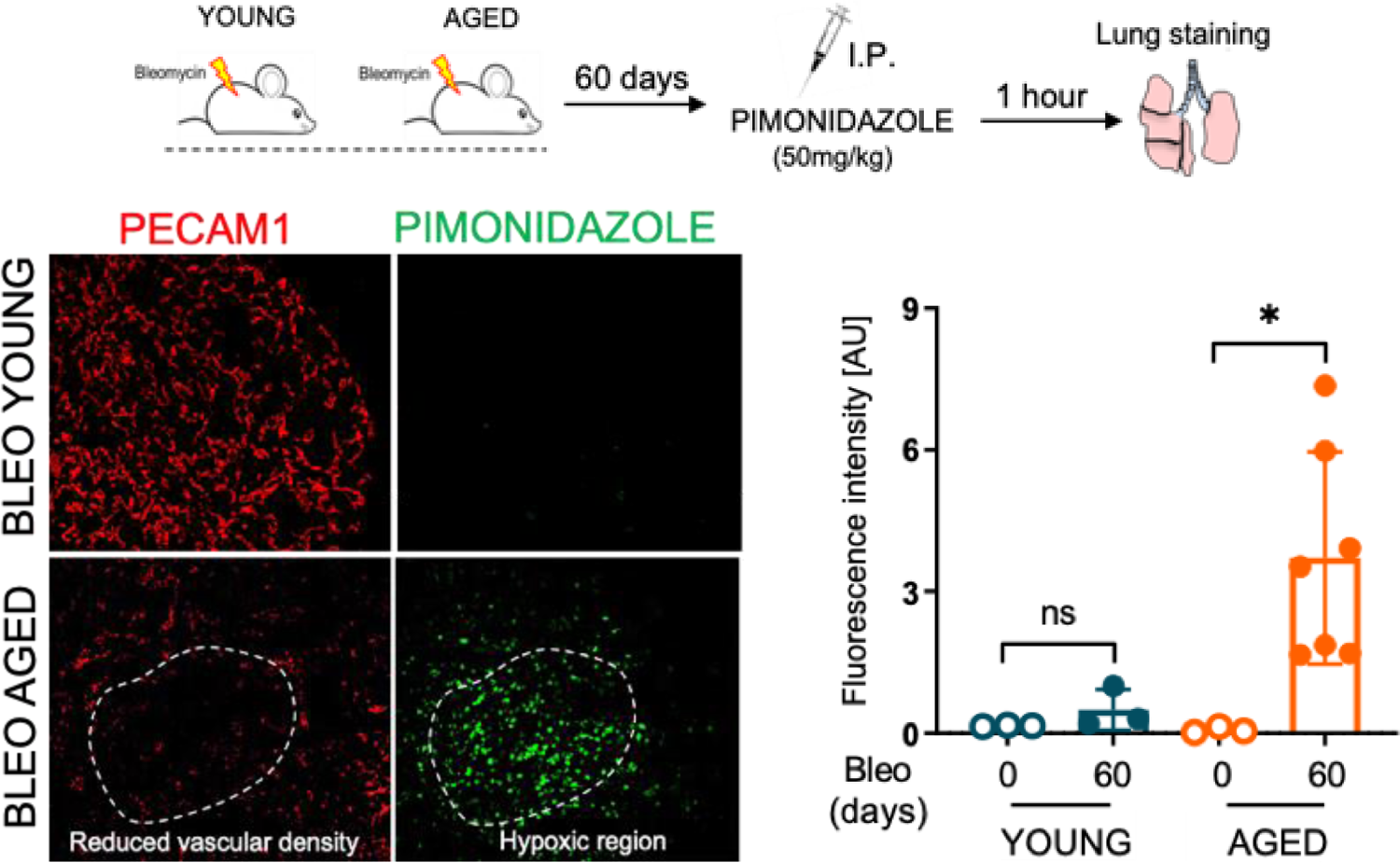
Elevated hypoxia in aged lungs compared to young ones following bleomycin-induced lung fibrosis. Hypoxia detection by Pimonidazole in young and aged mouse lungs at 60 days post bleomycin instillation. Immunofluorescence staining using antibodies against PECAM-1 shows reduced number of alveolar ECs in fibrotic aged lungs exhibiting elevated hypoxia compared to young one in which hypoxia levels were undetectable *p<0.05 (n≥3).

**Supplement Figure 5.**
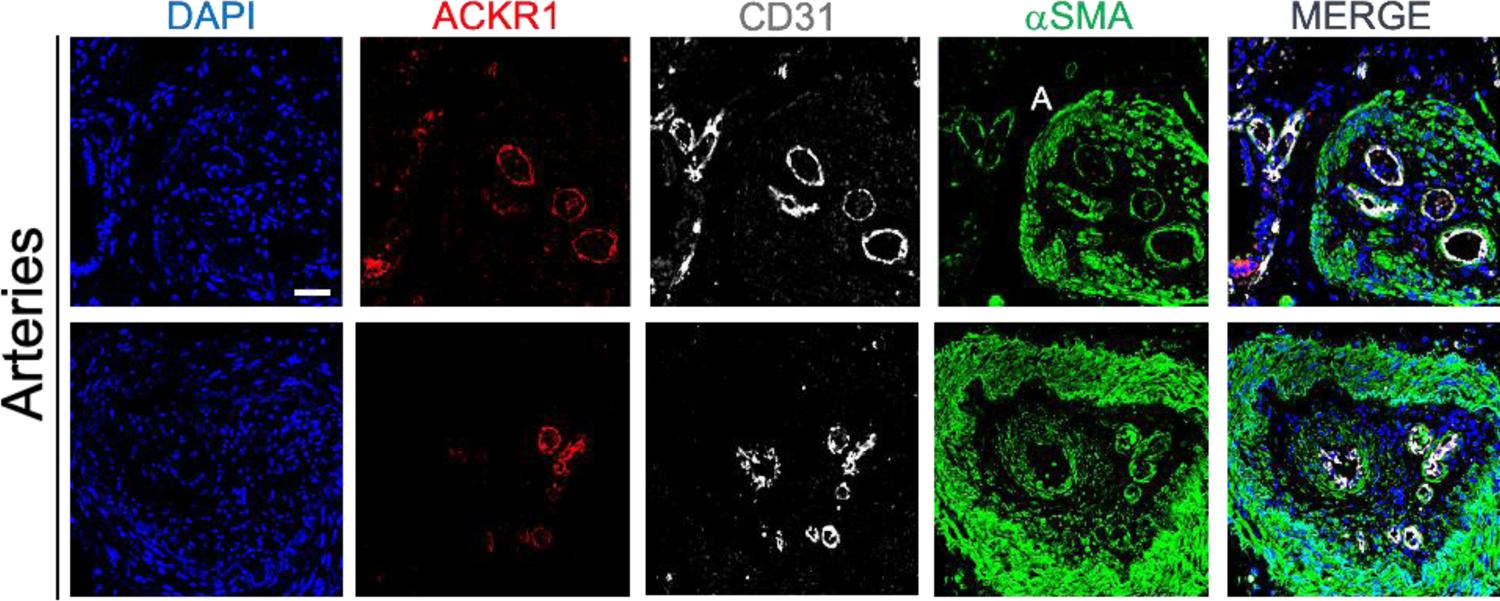
Vascular abnormalities involving ACKR1^+^ venous ECs in IPF lungs. ACKR1^+^ CD31^+^ venous ECs were found in the fibrotic intima of numerous arteries. Medial thickening and αSMA^+^ cells within the thickened intima were also observed in these abnormal vessels. Scale bar 50 µm.

